# KMT2D Regulates thymic Egress by Modulating Maturation and integrin Expression

**DOI:** 10.1101/2022.10.04.510662

**Authors:** Sarah J Potter, Li Zhang, Michael Kotliar, Yuehong Wu, Caitlin Schafer, Kurtis Stefan, Leandros Boukas, Dima Qu’d, Olaf Bodamer, Brittany N Simpson, Artem Barski, Andrew W Lindsley, Hans T Bjornsson

## Abstract

**Objective:** There is a clinical need to understand how dysregulated thymocyte development, caused by pathogenic variants in the gene encoding the histone-modifying enzyme, lysine methyltransferase 2D (*KMT2D*), contributes to immune dysfunction, including immune deficiency, autoimmunity, and lymphoproliferative sequela, and immune-driven mortality in individuals with Kabuki syndrome type 1 (KS1).

**Methods:** We studied peripheral T cells and thymocytes in both individuals with KS1 and murine constitutive and conditional targeted *Kmt2d* in T cells and hematopoietic lineages. KMT2D target genes, identified by RNA-sequencing of murine *Kmt2d*-knockout single-positive thymocytes, were validated with H3K4me3 ChIP-PCR and flow cytometry.

**Results:** Recent thymic emigrant (RTE) and naïve T cells were reduced, and memory and double-negative (DN)-T cells were expanded in human KS1 and murine models. *Kmt2d* loss led to Mature 1 CD8^+^-single positive (SP) thymocyte accumulation and a decrease in SP thymocyte egress licensing expression (normally associated with the Mature 2 phenotype). Splenomegaly is associated with hematopoietic-driven *Kmt2d* loss and brings to light potential overlapping phenotypes with lymphoproliferative syndromes. Finally, we identified a KMT2D-regulated cluster of integrins which likely mediates aspects of the T cell egression.

**Conclusions:** Single-positive thymocyte populations deficient in *Kmt2d* display less integrin, less maturation, and less egress licensing gene expression; thereby, altering the downstream peripheral T cell composition that contribute to the observed KS1-associated immune deficiency. T cell intrinsic *Kmt2d* loss increases the percentage of peripheral DNT cells potentially through dysregulated apoptotic signaling, while hematopoietic-driven *Kmt2d* loss predisposes to splenomegaly; therefore, loss of *Kmt2d* recapitulates several distinct features of lymphoproliferative syndromes.

## Introduction

Epigenetic dysregulation has emerged as a pathologic driver of many diseases, including primary immune deficiency. Kabuki syndrome type 1 (KS1) is an autosomal dominant, congenital, epigenetic disorder associated with heterozygous loss-of-function variants in the lysine methyltransferase 2D gene, *KMT2D* (also known as *MLL2/MLL4/ALR*). KS1 has a complex phenotype, including disruption of craniofacial, skeletal, cardiac, cognitive, and immune development. KS1 is frequently associated with specific forms of immune dysfunction, including humoral immunodeficiency, autoimmunity, and lymphoproliferation, which contribute to the morbidity seen in individuals with KS1 ^1–3^. A subset of these immune phenotypes can be explained through the known role of KMT2D in B cell survival/cell cycle regulation, differentiation, and peripheral tissue homing ^4,5^. For instance, KMT2D influences expression of its target gene integrin beta 7 (*ITGB7*) and the ITGA4-ITGB7 integrin heterodimer (also known as Lymphocyte Peyer’s Patch adhesion molecule-1 [LPAM1]) in mesenteric lymph node–derived lymphocytes ^6^. The contribution of T cell expressed KMT2D in KS1-associated immune deficiency (KSAID) pathogenesis and how *KMT2D* haploinsufficiency impacts the peripheral T lymphocyte population are currently poorly understood.

Recently, researchers have linked KMT2D to T cell dysfunction, indicating the potential contribution of abnormal T cells to KSAID. Specifically, 25 of 125 individuals (20%) with peripheral T-cell lymphomas not otherwise specified (PTCL-NOS) had *KMT2D* mutations. PTCL-NOS and healthy T cells can be distinguished by expression of cell cycle and adhesion genes that are dysregulated in PTCL-NOS ^7,8^. However, the contribution of T cells to the KS1 phenotype has remained unclear. Early immune phenotyping studies of individuals with KS1 suggested a decrease in the CD4^+^ memory compartment ^9^, but re-evaluation of these data have raised questions about whether the appropriate reference ranges were used ^10^. Further studies also investigated KMT2D’s role in early T cell development via characterizing long-range chromatin interactions (enhancers) in thymocytes and naïve T cells and determined that *Kmt2d* loss contributed to a reduction in thymic CD4^+^ T regulatory cells, and in peripheral CD4^+^ and CD8^+^ T cells ^11^.

Herein, we show that KMT2D plays a crucial role in thymocyte development, egress, and peripheral expansion through regulating integrin expression of ITGAE, ITGAL, and ITGB7. Additionally, given that *Kmt2d* deficiency leads to increased double-negative T (DNT) cells and splenomegaly, we demonstrate that a key *Kmt2d* function in hematopoietic cells is to suppress pathogenic lymphoproliferation through both T cell-intrinsic and -extrinsic mechanisms.

## Results

### Human recent thymic emigrants are decreased in individuals with molecularly confirmed KS1

To analyze T lymphocyte disruption in KS1, we recruited a group of individuals (n = 16) with diverse *KMT2D* genetic variants deemed likely pathogenic/pathogenic by American College of Medical Genetics and Genomics criteria and diagnosed with KS1 by a clinical geneticist (**Figure 1a/Table 1**). Upon flow cytometric analysis of peripheral blood cells, we found that CD3^+^ T cells and CD4^+^ and CD8^+^ T cell subpopulations were within the age-matched clinical reference ranges (**Figure 1b-d**). Interestingly, DNT cells were above the reference threshold in all, but one patient (**Figure 1e-f**). Upon further analysis of the CD4^+^/CD8^+^ subpopulations, we observed individuals with KS1 have a significantly lower percentage of naïve CD4^+^ T cells and a corresponding expansion of memory CD4^+^ T cells; however, the corresponding CD8^+^ T cell compartments were within the clinical reference range (**Figure 1g-h/ Supplementary figure 1a-b**). Furthermore, individuals with KS1 had a significant reduction in the percentage of CD4^+^CD31^+^ recent thymic emigrants (RTE; **Figure 1i**). Given these consistent T cell abnormalities in human individuals with KS1, we further explored the T cell compartment in relevant murine models.

**Figure 1.**
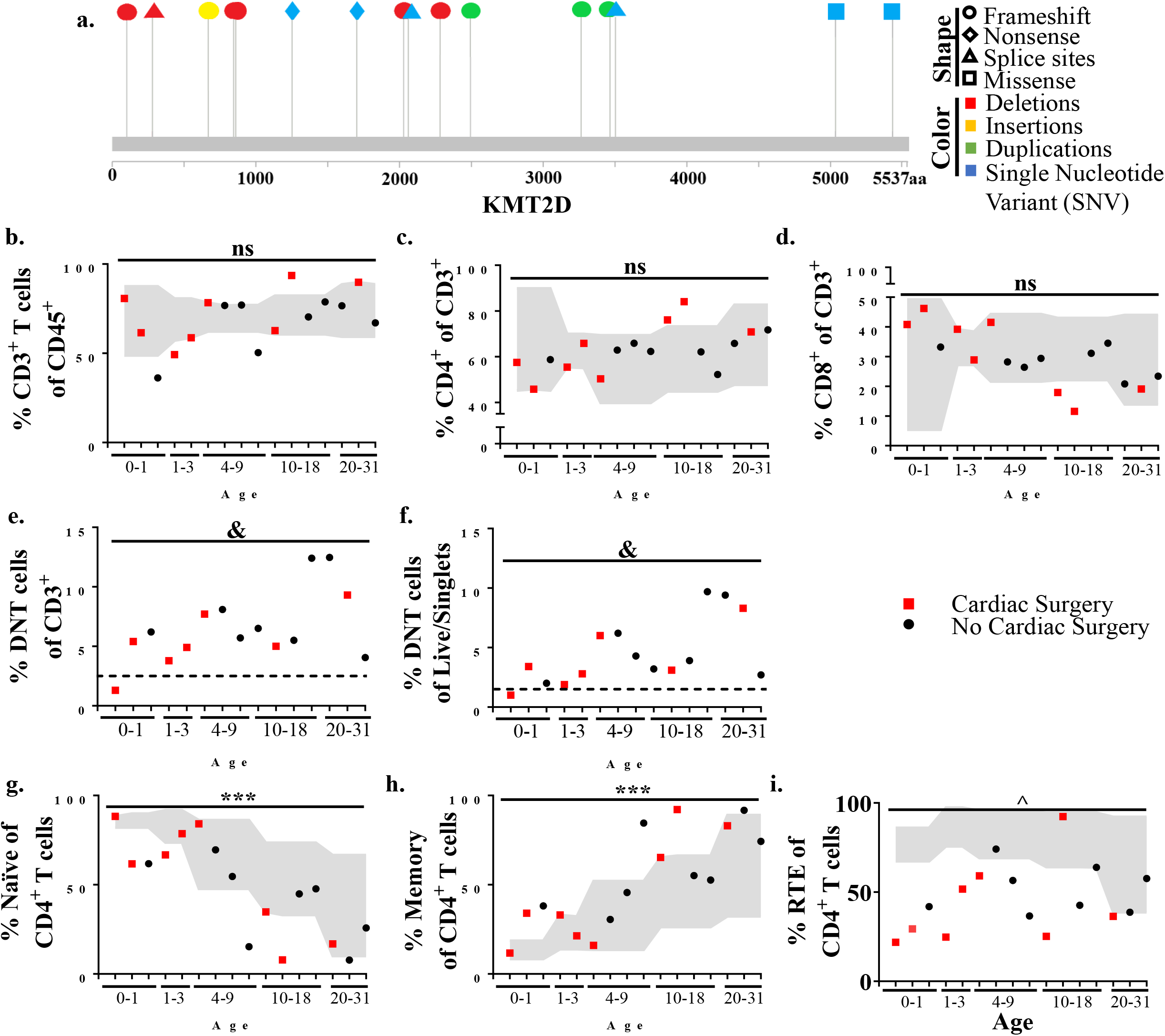
Individuals with KS1 display decreased T cell counts, naïve CD4^+^ T cells, and RTE CD4^+^ T cells, and increased memory CD4^+^ T cells. (**a**) Individual variants found in KS1 cohort (n = 16; ages 0 – 31). Lolliplot indicates protein amino acid location (via stick location), types of variant (i.e. shape: squares: missense, circles: frameshifts, triangles: splice sites, diamonds: nonsense), and changes at the nucleotide level (i.e. color: deletions: red, insertions: yellow, duplications: green, and single nucleotide variant (SNV): blue). (**b-i**) Data from clinical flow cytometry (markers represent separate individuals). Cardiac surgery (red square; non-surgery: black circle), required for (or performed on) many individuals with KS1, can remove and/or damage the thymus. (**b**) Percentage of peripheral T CD45^+^CD3^+^ T cells of total blood cells. (**c**) Percentage of CD4^+^ T cells of CD3^+^ T cells. (**d**) Percentage of CD8^+^ T cells of CD3^+^ T cells. (**e-f**) DNT cells (CD3^+^CD4^-^CD8^-^) as a percent of CD3^+^ cells (**e**) or as a percent of CD45^+^ cells (**f**). Non age-matched reference range above 2.5% or 1.5% DNTs in (**e**) and (**f**) populations, respectively, is considered variant (threshold indicated by dotted line). (**g**) Naive CD4^+^ T cells (CD45RA^+^CD27^+^) as a percent of CD4^+^ T cells. (**h**) Memory CD4^+^ T cells (CD45RA^-^) as a percent of CD4^+^ T cells. (**i**) Recent thymic emigrants (RTE; CD27^+^CD45RA^+^CD31^+^) as a percent of CD4^+^ naïve T cells. (**b**-**d, g**-**i**) Values found within grey region represents reference population range (2.5% - 97.5%) for the healthy age-matched individuals. Significance was determined using the binomial test, with the null hypothesis being that individuals with KS1 have a same probability of falling outside the reference range equal to 5%, that is, the same as healthy age-matched individuals. The resulting *P*-values were subsequently adjusted for multiple testing with the Bonferroni method and noted by asterisks: *P*-value > 0.05 (ns), *P*-value < 0.001 (***), *P*-value < 1e^-9^ (^), and *P*-value < 1e^-14^ (&). Specifically, in (**b**), CD3^+^ significance were calculated for values below the standard range only (one-tailed binomial test).

**Table-1.**
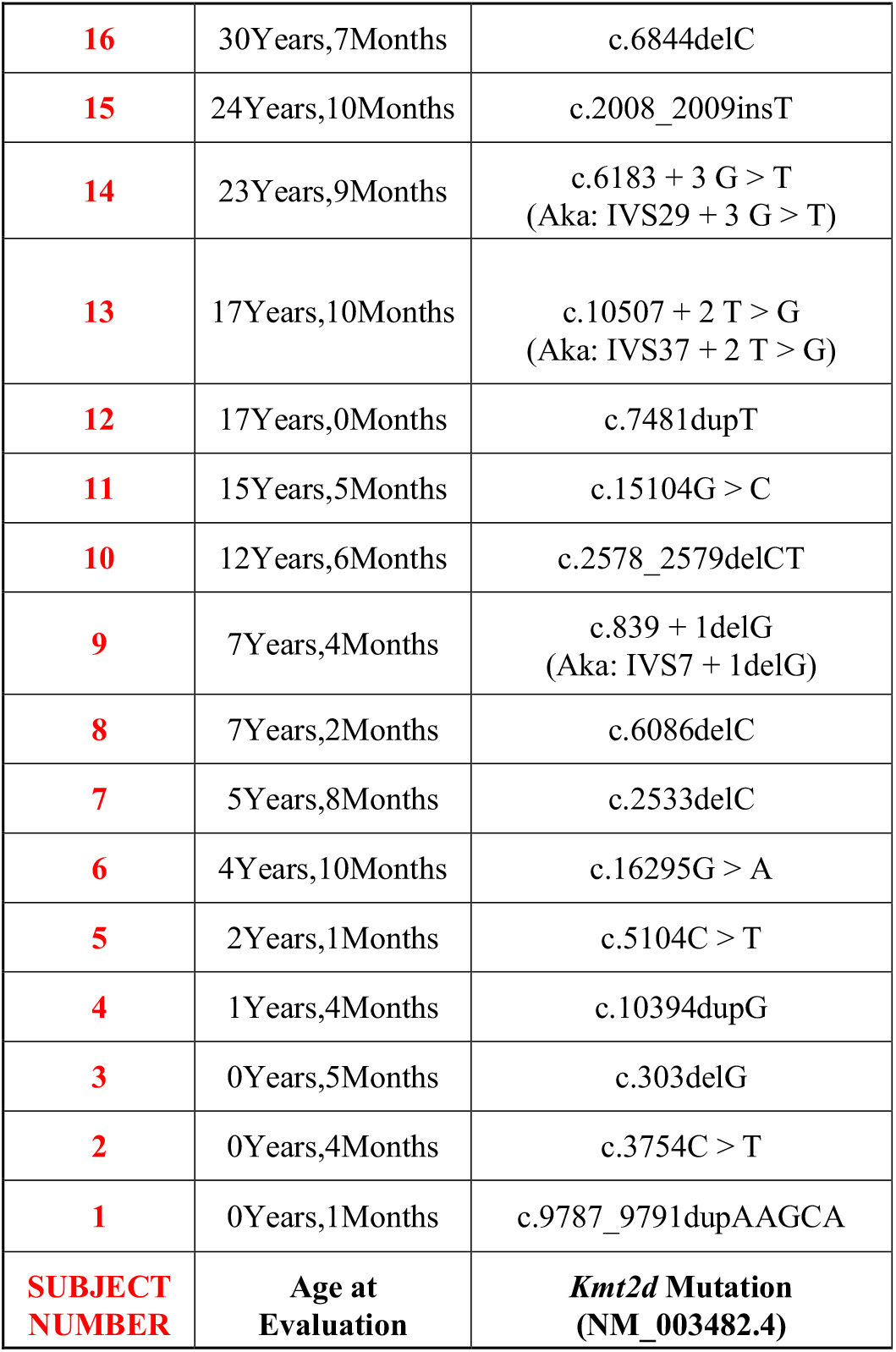
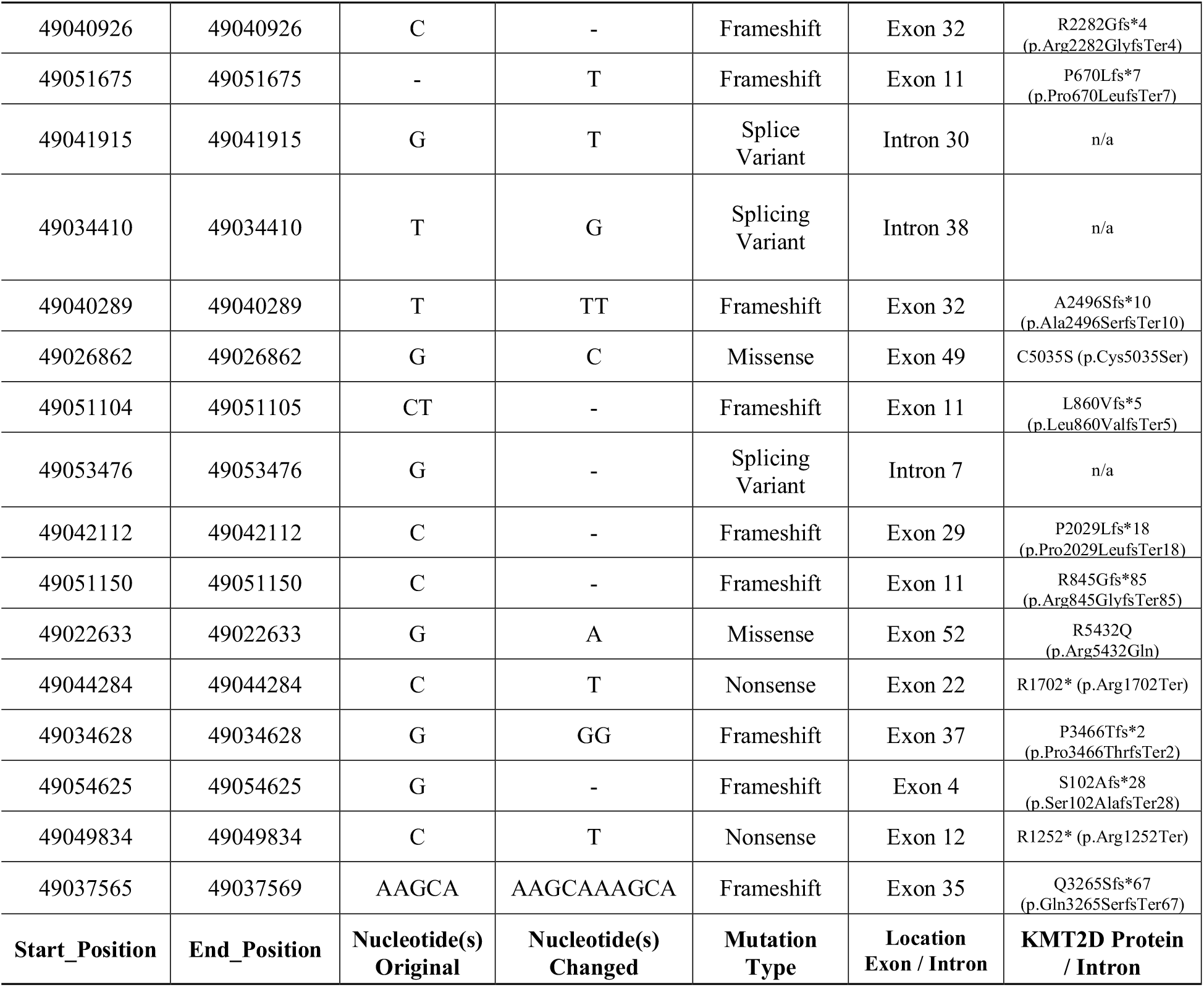
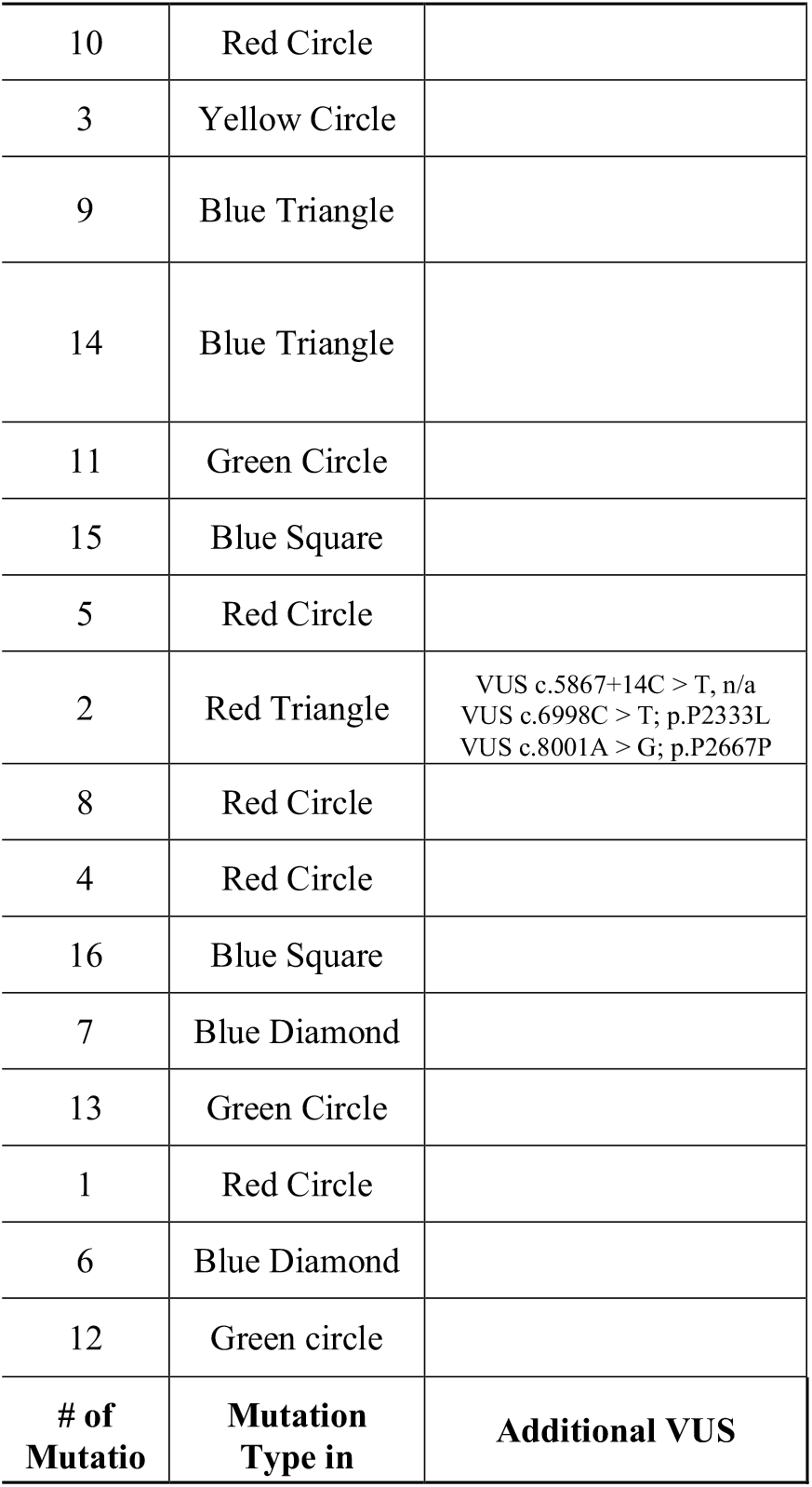
De-identified information collected from individuals with KS1. Data from this information corresponding to Figure-1/ Supplementary figure 1. Each column represents an individual. **Figure-1** graphs are assembled in ascending order (Row 1) based off the individuals’ age at the time of lymphocyte evaluation (Row 2). The following information are provided: location of the KMT2D variation and the type of change (single-nucleotide variation, duplication, insertion, deletion, etc; Row 3); information, such as start position (Row 4) and end position (Row 5) as located on the chromosome 12, nucleotide(s) of the reference genome (Row 6) and what the variation changes in the individual (Row 7) were required to generate the lolliplot (Rows 4-7); VarSome determination of the variation type (missense, nonsense, frameshift, and splice variant; Row 8); location where the variation takes place (within what *KMT2D* exon/intron; Row 9); VarSome KMT2D protein amino acid prediction (Row 10); location of the amino acid change on the lolliplot (Row 11) and the type (noted by shape/color; Row 12); and known locations of variants of uncertain significance (VUS) for the individual are indicated (Rows 13-15). Abbreviations found in Row 3: † insertion (ins), ‡ deletion (del), § duplication (dup), ¶ single nucleotide change (>).

### Peripheral T cell changes in Kmt2d-deficient mice recapitulate human phenotype

To understand whether peripheral T lymphocytes are influenced by the loss of *Kmt2d*, we analyzed splenic T cell composition in an established, constitutive haploinsufficient mouse model, herein called *Kmt2d^+/βgeo^* mice, and two conditional T cell–specific *Kmt2d* knockouts, driven by *CD4-Cre* or *Lck-Cre*^Mar^, that initially recombine at distinct late DN thymocyte stages (**Supplementary figure 2)**. Also, in limited cases, we have employed a conditional hematopoietic-activated *Kmt2d* knockout driven by *Vav1-iCre* ^12^. Similar to the human KS1 cohort, *Kmt2d^+/βgeo^* (haploinsufficient) mice had no significant changes in CD3^+^ T cell percentages from littermate controls, whereas all three conditional *Kmt2d*-knockout models (*CD4*-Cre,*Lck*-Cre^Mar^, *Vav1*-iCre) had significant decreases in CD3^+^ T cells **Figure 2a/ Supplementary figure 3a**). Across all *Kmt2d*-deficient models a significant reduction in the CD4^+^ subpopulation percentage was noted and correspondingly the DNT cell subpopulation percentage increased, while CD8^+^ subpopulation percentage remained similar to control levels; resulting in skewed CD4^+^/CD8^+^ ratios toward CD8^+^ (**Figure 2b/ Supplementary figure 3c**). Within *Kmt2d*-deficient animals, overall CD3^+^CD4^+^ and CD3^+^CD8^+^ populations were decreased, while DNTs were increased (**Supplementary figure 3a-b**). Furthermore, we observed reduced naïve and enhanced memory CD8^+^ population in *Kmt2d*-targeted mouse models (**Figure 2c**) that mirror the CD4^+^ naïve/memory alterations seen in individuals with KS1.

**Figure 2.**
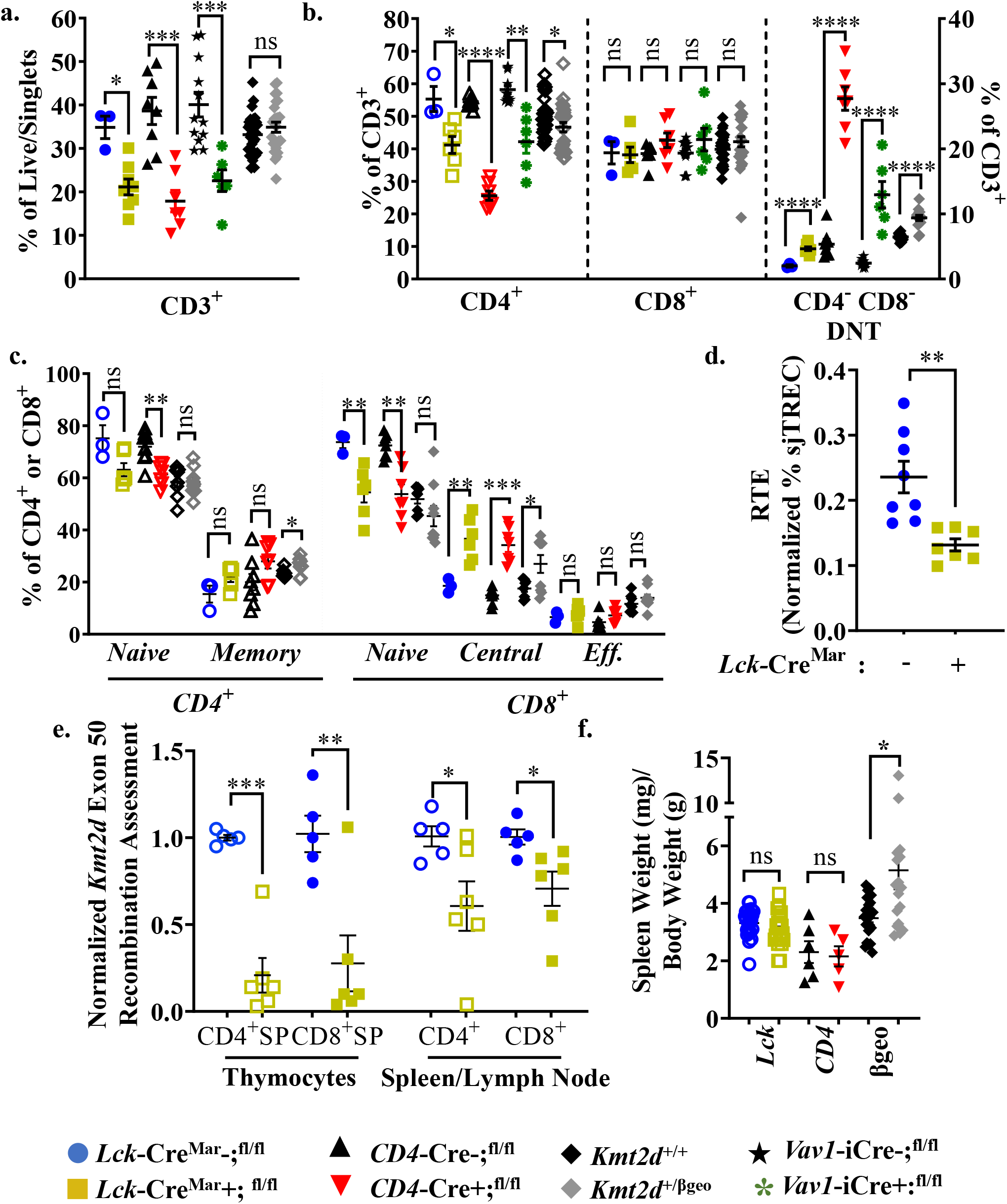
*Kmt2d*-deficient peripheral T cells demonstrate decreased total T cells, naïve T cells, and recent thymic emigrant T cells, but increased memory phenotypes. (**a**) Flow cytometry of peripheral CD3^+^ as percent of live/singlet cells. (**b**) Peripheral CD4^+^ (open, left), CD8^+^ (closed, middle), and DNT (closed, right with scale on right) T cell populations of total CD3^+^ shifts. (**c**) Peripheral splenic naïve and memory CD4^+^ (open) and CD8^+^ (closed) T cell population dynamics (based on CD62L and CD44 expression). (**d**) Total peripheral blood recent thymic emigrant changes shown through sjTREC PCR values normalized to CD3^+^ percent in *Lck*-Cre^Mar^; *Kmt2d*^-SET-fl/fl^. (**e**) Genomic DNA recombination assessment of exon 50 (SET domain), which is cleaved during Cre-recombination at *Loxp3* sites, in thymocytes and peripheral spleen/lymph node CD4^+^ (open) and CD8^+^ (closed) T cells of *Lck*-Cre^Mar^; *Kmt2*^-SET-fl/fl^ and littermate controls. Calculation plotted is 2^-(Mutant [CT of *Kmt2d*-Exon50 / CT of *Kmt2d*-Total R20] – Control [CT of *Kmt2d*-Exon50 / CT of *Kmt2d*-Total R20])^, where the knockouts are normalized to all *Kmt2d* and then normalized to the average of the cell specific *Lck*-Cre^Mar-^; *Kmt2d*^-SET-fl/fl^ control; see schematic in **Supplementary figure 2**. (**f**) Spleen weight (mg) normalized to total weight (g). (**a-e**) The following mouse lines are displayed as splenic cells (or as listed: thymocytes [**e**] or peripheral blood [**d**]) from individual mice (n = 3 - 30) on the graphs: *Kmt2d^+/βgeo^*(^+/+^ black diamonds/^+/βgeo^ grey diamonds), *Lck*-Cre^Mar^; *Kmt2d*^-SET-fl/fl^ (Cre^-^ blue circles/Cre^+^ yellow squares), *CD4*-Cre; *Kmt2d*^-SET-fl/fl^ (Cre^-^ black up-right triangles/Cre^+^ red inverted triangles), and peripheral blood *Vav1*-iCre; *Kmt2d*^-SET-fl/fl^ (Cre^-^ black stars/Cre^+^ green asterisks). Each dataset displays black lines, which represent mean ± SEM. Significance labeled on graphs were determined using a parametric, unpaired, Welch’s corrected *t-test* with *P*-values noted by asterisks: *P*-value > 0.05 (ns), *P*-value < 0.05 (*), *P*-value < 0.01 (**), *P*-value < 0.001 (***), and *P*-value < 0.0001 (****).

To evaluate thymic egress in *Kmt2d* knockout T cells, we employed the sjTREC assay, which quantifies RTE cells at a population level ^13^. Specifically, T cell receptor (TCR) recombination generates excised V(D)J loops, whose concentration is diluted upon each subsequent round of cell division. We observed a significant decrease in CD3^+^ normalized sjTREC signal in *Kmt2d*-knockout cells compared to controls, but not in *Kmt2d*-heterozygous cells (**Figure 2d/ Supplementary figure 3d)**. The decreased sjTREC signal could indicate that the peripheral *Kmt2d* knockout T cells either have: 1) undergone more cellular divisions than matched control cells; and/or 2) as a population contain a smaller fraction of RTE than *Kmt2d*-sufficient T cells; or 3) have been subject to a combination of both excessive proliferation and reduced thymic egress. Overall, the observed decrease in sjTREC signal is consistent with the previously identified reduced naïve cells and enhanced peripheral expansion of memory T cells in *Kmt2d* knockout models.

To further characterize the role of KMT2D in thymic egress, we evaluated *Kmt2d* locus targeting (i.e. knockout) efficiency via quantification of Cre-recombinase cleavage of exons 50/51 (SET-domain) in the *Kmt2d*^SET-fl/fl^ mice, in both thymocytes and peripheral T cells (**Supplementary figure 2**). *Kmt2d*-knockout peripheral T cells showed decreased SET deletion than their SP thymocyte precursors (**Figure 2e**). The over-representation of peripheral *Kmt2d*-sufficient cells, which have “escaped” CRE recombination despite expected levels of thymic recombination, suggests that the complete loss of *Kmt2d* is detrimental for thymic egress, proliferation, and/or viability. Interestingly, *Kmt2d*-heterozygous thymocytes and peripheral cells have a similar *Kmt2d* recombination efficiency (**Supplementary figure 3e**). This demonstrates that one allele of *Kmt2d* is sufficient to sustain T cell egress.

Lastly, in addition to the other peripheral phenotypes, we observed splenomegaly in whole-body haploinsufficient and hematopoietic (*Vav1*)-driven, but not with T cell (*Lck/CD4*)-driven KMT2D deletions. Therefore, splenomegaly appears to be induced through T cell extrinsic mechanisms (**Figure 2f/ Supplementary figure 3f-g**).

### Altered thymus weight and thymocyte populations within constitutive haploinsufficient and conditional homozygous targeting of Kmt2d

To evaluate the cause of the RTE deficiency, we examined thymic development in mice with conditional T cell-specific loss of *Kmt2d*. First, we analyzed murine thymus weight and found no significant change in *Kmt2d^+/βgeo^* mice, nor *CD4*-Cre-driven *Kmt2d* knockouts; however, the T cell–specific *Lck*-driven *Kmt2d*-knockout led to a significantly increased thymus weight (**Figure 3a**/ **Supplementary figure 4a**). Why thymic the hypertrophy phenotype differed between the two T cell-specific *Kmt2d* knockout models in unclear; however, one possible explanation is the timing of the Cre recombinase expression differs slightly between models (with *Lck*-Cre^Mar^ turning on DN3 compared to DN4 for *CD4*-Cre) ^12^. To elucidate the stages in which *Kmt2d* plays a role during thymocyte development, we comprehensively analyzed thymocyte populations (CD4^-^CD8^-^DN, CD4^+^CD8^+^DP, CD8^+^SP and CD4^+^SP) across developmental time. Strikingly, *Kmt2d* loss significantly increased the percentage of CD8^+^SP thymocytes (of live/single cells) in both *Kmt2d*-knockout and the *Kmt2d*-haploinsufficent models (**Figure 3b**).

**Figure 3.**
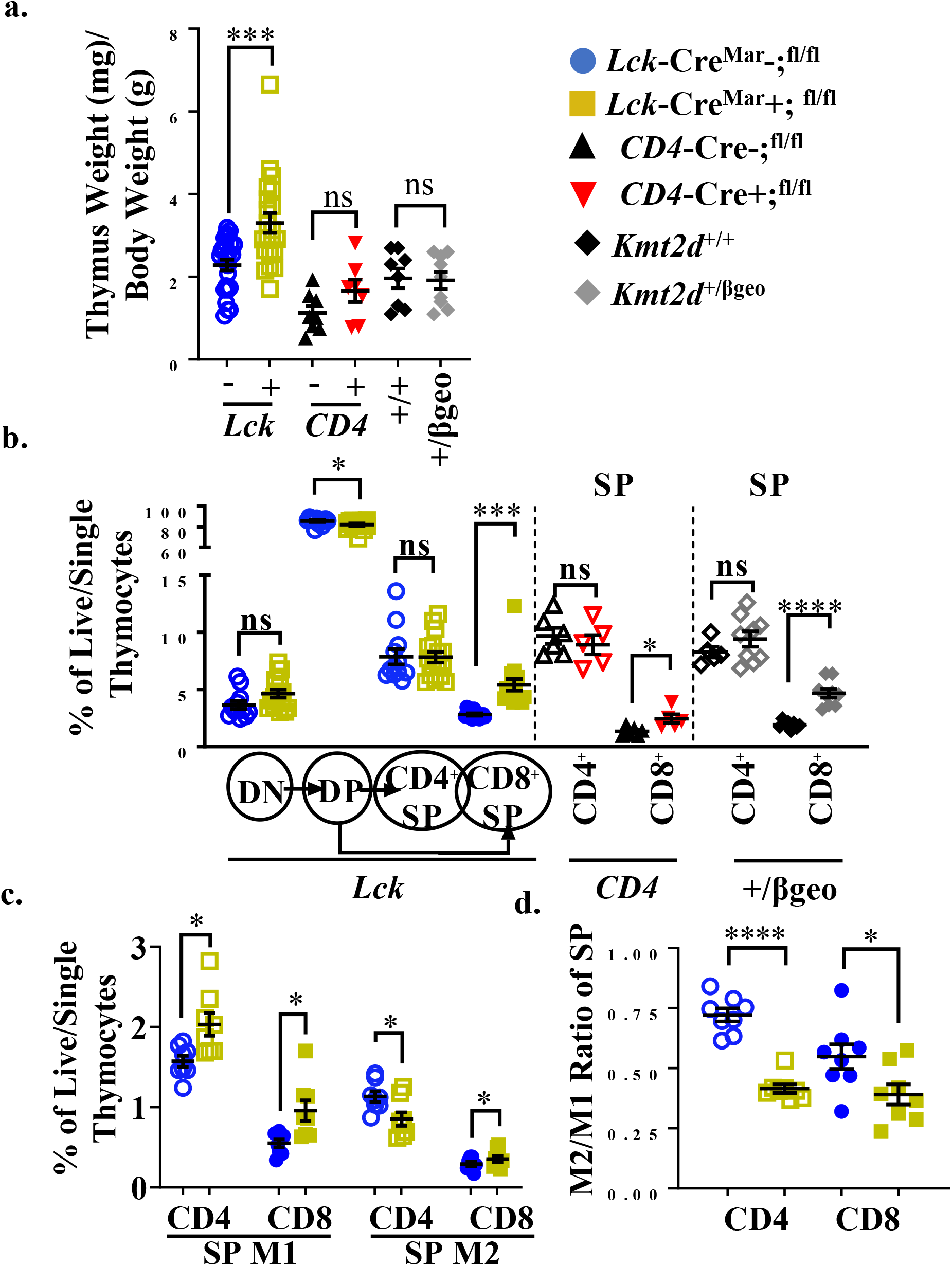
Accumulation of mature 1 CD8^+^SP cells in *Kmt2d-deficient* mice. (**a**) Organ weight of thymus (milligrams) normalized to body weight (grams). (**b**) Flow cytometric single cell analysis of thymocyte development populations as indicated: double-negative (DN), double-positive (DP), and single-positives (CD4^+^SP, CD8^+^SP) as a percent of total live/single thymocytes, of *Lck^-^*Cre^Mar^; *Kmt2d*^-SET-fl/fl^ thymocyte populations (left) and CD4^+^SP and CD8^+^SP from *CD4*- Cre; *Kmt2d*^-SET-fl/fl^ (middle), then *Kmt2d^+/βgeo^* (right) separated by dotted lines. CD8^+^SP are significantly reduced in all models. (**c**/**d**) Thymic SP maturation, progressing from Mature 1 MHCI^+^CD69^+^CCR7^+^TCRβ^+^ (M1; left) to MHCI^+^CD69^-^CCR7^+^TCRβ^+^ Mature 2 (M2; right) of *Lck*-Cre^Mar^; *Kmt2d*^-SET-fl/fl^ CD4^+^SP (open) and CD8^+^SP (closed) thymocytes as a percent of total thymocytes (**c**) by flow cytometry or by ratio of percent of total thymocytes M2/ percent of total thymocytes M1 (**d**). The following mouse lines are displayed as cells from individual mice (n = 5 - 15) on the graphs: *Kmt2d^+/βgeo^* (^+/+^ black diamonds/^+/*βgeo*^ grey diamonds), *Lck*-Cre^Mar^; *Kmt2d*^-SET-fl/fl^ (Cre^-^ blue circles/Cre^+^ yellow squares), and *CD4-Cre; Kmt2d*^-SET-fl/fl^ (Cre^-^ black up-right triangles/Cre^+^ red inverted triangles). Each dataset displays black lines, which represent mean ± SEM. Significance is listed on graph based on *P*-value > 0.05 (ns), *P*-value < 0.05 (*), *P*-value < 0.01 (**), *P*-value < 0.001 (***), and *P*-value < 0.0001 (****).

Thymocyte maturation occurs concurrently with negative selection (contributing to the CD4^+^ T regulatory cell pool and removal of auto-reactive CD4^+^ and CD8^+^ T cells) as a multi-stage process. It proceeds from semi-mature cells into the Mature 1 (M1)-SP thymocytes, which are not completely “licensed” for egress, and lastly into licensed for egress Mature 2 (M2)-SP thymocytes. *Kmt2d*-knockout thymocytes have a significantly increased percentage of M1-SP (**Figure 3c**). Furthermore, within T cell-specific *Kmt2d* knockout mice, CD4^+^SP thymocytes experience an increase in M1-SP thymocytes, but a decrease in M2-SP thymocytes compared to their control littermates; therefore, not contributing to changes in the overall CD4^+^SP population (**Figure 3b-d**). In contrast, CD8^+^SP demonstrate an increase in both M1-SP and M2-SP thymocyte populations (**Figure 3b-d**) leading to an increase in the overall CD8^+^SP population. Furthermore, the CD44 cleavage during SP maturation required for egress ^14^ did not occur as frequently within *Kmt2d*-deficient SP cells (**Supplementary figure 4b**). Together, these results suggest *Kmt2d* loss causes a block in M1-M2-SP thymocytes transition, which may impact egression.

### Kmt2d-regulated genes during thymocyte differentiation

KMT2D is a co-activator that regulates gene activation through histone H3K4 methylation on promoters and enhancers; therefore, we expect that the genes with reduced RNA expression in *Kmt2d*-knockout compared to control cells may be directly regulated by *Kmt2d*, whereas genes with increased expression are likely secondary effects, i.e. bystander or negatively-regulated downstream genes. To identify *Kmt2d*-regulated thymocyte genes and downstream pathways, we performed RNA-sequencing (RNA-seq) on *Kmt2d*-knockout CD4^+^SP and/or CD8^+^SP thymocytes from either T cell–specific (*Lck*-Cre^Mar^) or hematopoietic stem cell-activated (*Vav1*-iCre) models (**Supplementary figure 5-9**). *Kmt2d* loss yielded down-regulated genes with top biological functions relating to activation, cell adhesion, and GTPases (**Figure 4a**).

**Figure 4.**
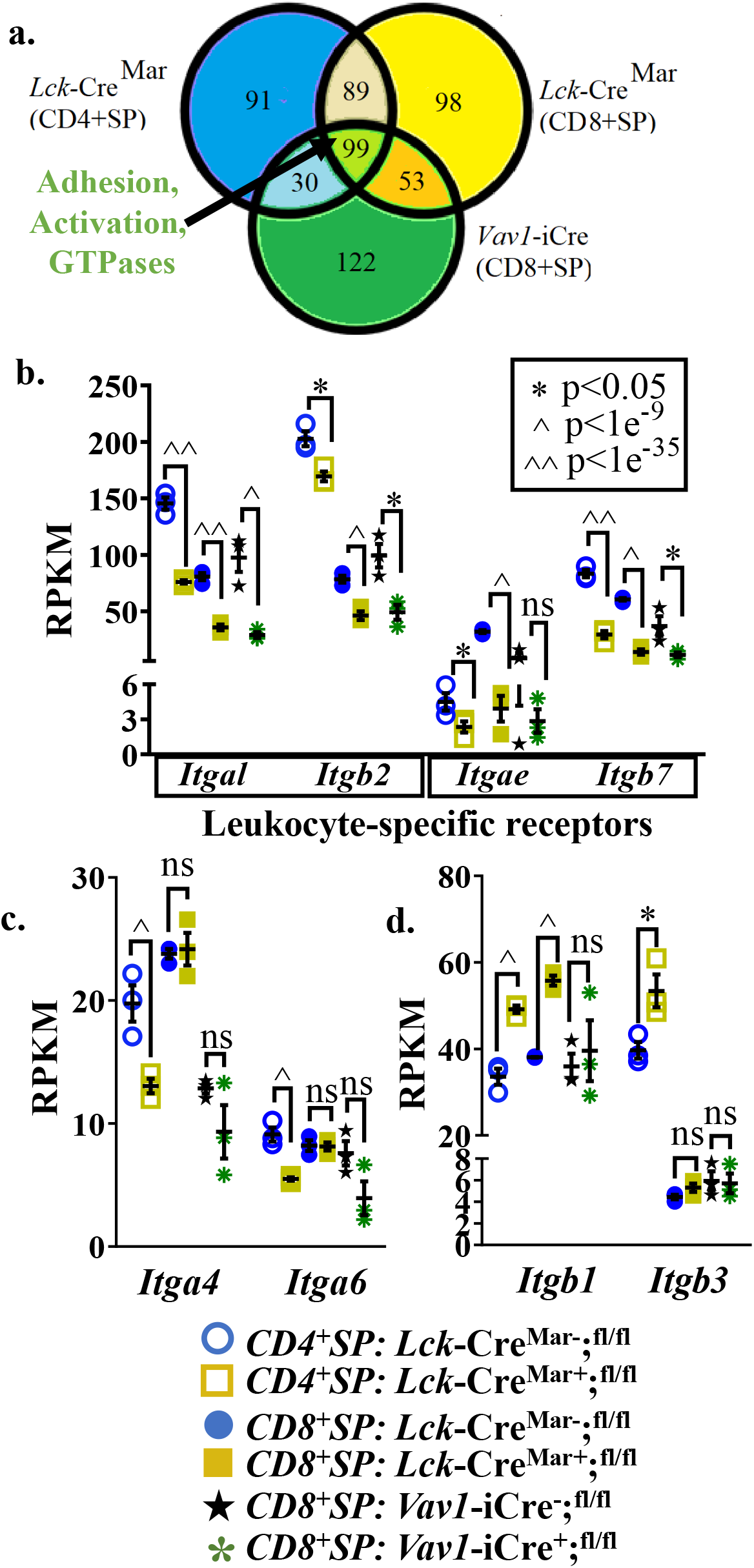
Expression analysis in *Kmt2d*-knockout mice reveals systematic disruption of leukocyte-specific receptors. (**a**) Potential *Kmt2d* target genes estimated by selection of down-regulated genes in the *Kmt2d*-knockout SP thymocytes compared to controls SP thymocytes as *Kmt2d* normally regulates gene expression. Venn diagram demonstrates overlapping *Kmt2d*-knockout down-regulated filtered gene lists (Log2 Fold Change ≥ 0.59; adjusted *P*-value ≤ 0.1, RPKM ≥ 5 in at least one condition) between *Kmt2d* CD8^+^SP/CD4^+^SP (**a**) as indicated from RNA-seq DESeq2 from *Vav1-iCre* (green) and *Lck*-Cre^Mar^ (yellow) knockout CD8^+^SP cells and CD4^+^SP *Lck*-Cre^Mar^ knockouts (blue). The resulting shared 99 down-regulated genes in the middle green oval/triangle have the top GO categories in activation, adhesion, and GTPases. (**b**-**c**) RPKM of leukocyte-specific receptor integrin (**b**) and non-leukocyte specific integrins (**c**) that are down-regulated in *Kmt2d* knockouts and up-regulated non-leukocyte specific integrins (**d**). (**b**-**d**) The following mouse lines are displayed as cells from individual mice (n = 3) on the graphs: *Kml2d^+/βgeo^* (^+/+^ black diamonds/^+/*βgeo*^ grey diamonds), *Lck*-Cre^Mar^; *Kmt2d*^-SET-fl/fl^ (Cre^-^ blue circles/Cre^+^ yellow squares), and *CD4-Cre; Kmt2d*^-SET-fl/fl^ (Cre^-^ black up-right triangles/Cre^+^ red inverted triangles). Each dataset displays black lines, which represent mean ± SEM. Significance (adjusted *P*-values are generated from DESeq2 analysis *P*-value > 0.05 (ns), *P*-value < 0.05 (*), *P*-value < 1e^-9^ (^), and *P*-value < 1e^-35^ (^^).

Given our prior report of *Kmt2d* regulating *Itgb7* expression ^6^ and the biological function categories highlighting adhesion, we systematically focused on integrin molecules. Both *Kmt2d*-knockout and littermate control CD8^+^SP and CD4^+^SP thymocytes replicated the previously described expression of: *Itga4, Itga5, Itga6, Itgae* (only CD8^+^SP)*, Itgal, Itgav, Itgb1, Itg2, Itgb3*. ^15^ Although expressed, *Kmt2d* knockouts compared to controls had significantly decreased expression of *Itga4, Itga6* and in leukocyte-specific dimerizing partner integrin pairs, such as *Itgae/Itgb7* and *Itgal/Itgb2*, but increased *Itgb1* and *Itgb3* expression (**Figure 4b-d**).

As M1-SP thymocyte population accumulation and alterations of integrin gene expression were observed in *Kmt2d*-knockout SP thymocytes, we wanted to determine whether the FOXO1-driven egress licensing pathway was defective in SP thymocytes. Reportedly, FOXO1 is only expressed in the most mature SP cells (CD69^-^ and no longer receiving negative selection TCR signaling) and binds *Klf2* ^16–18^. Subsequently KLF2 binds a plethora of homing genes (*S1pr1* [encodes sphingosine-1-phosphate receptor 1, the receptor for S1P1], *Itgb7, Ccr7, Sell* [encodes CD62L]) required for egress. In *Kmt2d*-knockout CD4^+^SP and CD8^+^SP thymocytes had significantly reduced expression of *Foxo1, Klf2, Itgb7*, and *Ccr7*, but not *S1pr1* (**Figure 5**). Next, we evaluated these integrin and licensing gene targets of KMT2D at the chromatin regulation level.

**Figure 5.**
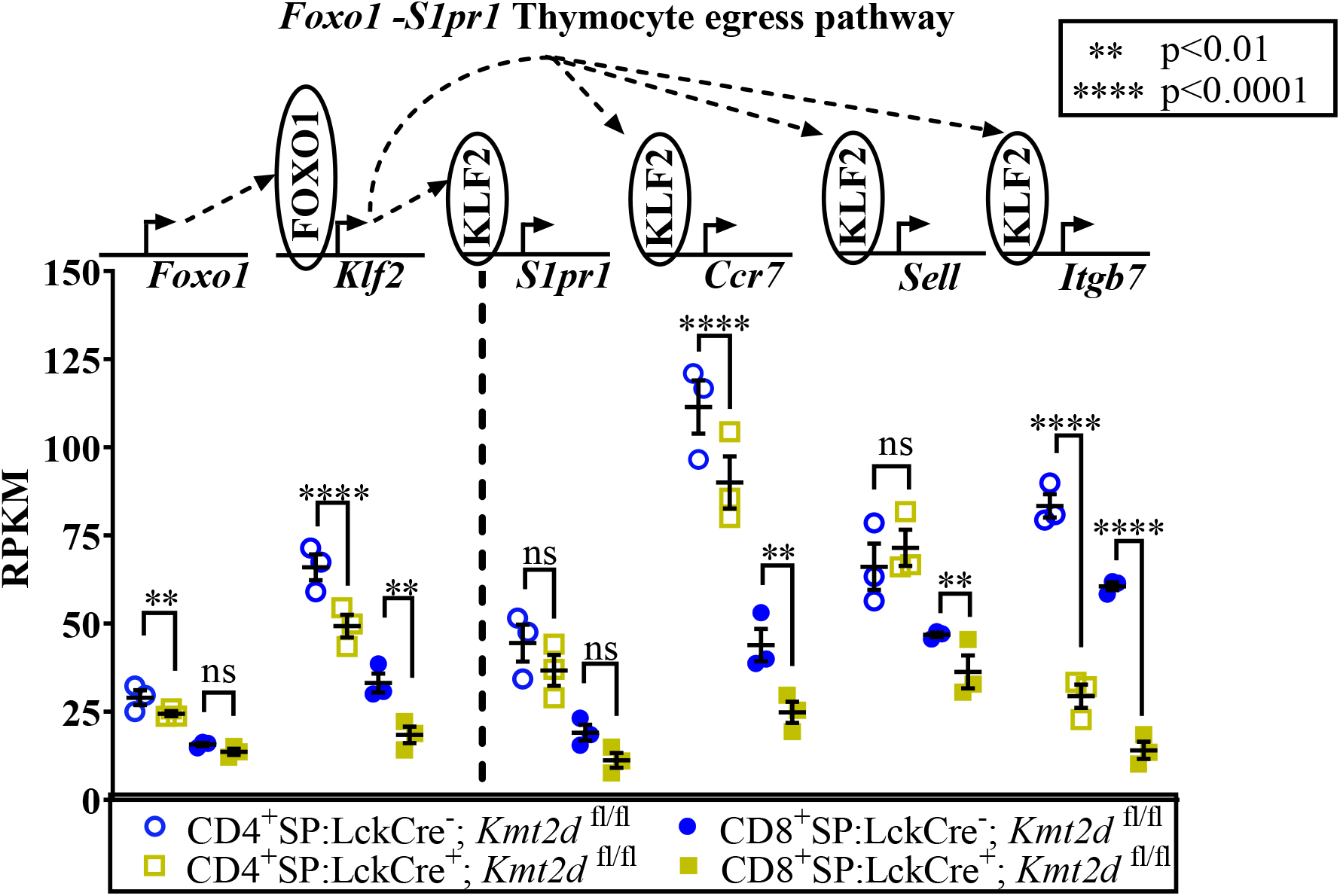
Disruption of the entire egress licensing pathway (*Foxo1-S1pr1* expression) in *Kmt2d* knockout SP cells. Individual RPKM of CD4^+^SP (open) and CD8^+^SP (closed) *Lck*-Cre^Mar^; *Kmt2d*^-SET-fl/fl^ (Cre^-^ blue circle/Cre ^+^ yellow square) from RNA-seq of the *Foxo1-Slpr1* pathway. Relevant genes displayed on the top of each dataset. Each dataset displays individual mice (n = 3; each marker) and also black lines, which represent mean ± SEM. Pathway depicted above graph: FOXO1 protein binds to promote gene expression of *Klf2*, fueling KLF2 protein expression, which binds to 4 downstream genes inducing increased RNA expression of the said genes. DESeq2 adjusted *P*-values listed on the graph indicate significance based on *P*-value > 0.05 (ns), *P*-value < 0.01 (**), and *P*-value < 0.0001 (****).

#### A subset of differentially expressed genes appear under direct KMT2D control

To examine whether KMT2D directly regulates SP thymocyte egress licensing/migration-associated genes, *Klf2, Itgal*, and *Itgb7*, we utilized published KMT2D chromatin immunoprecipitation-sequencing (ChIP-Seq) data (GEO: GSE69162) to identify KMT2D peaks present in putative cis-regulatory regions of these genes within CD4^+^ T cells (**Figure 6a**). Next, we performed ChIP-PCR to quantify levels of H3K4me3 at these cis-regulatory regions. We found significantly decreased H3K4me3 levels at these cis-regulatory regions, which suggest direct KMT2D-catalyzed chromatin modification; in contrast, the control *Gapdh* promoter region showed no such effect (**Figure 6b**).

**Figure 6.**
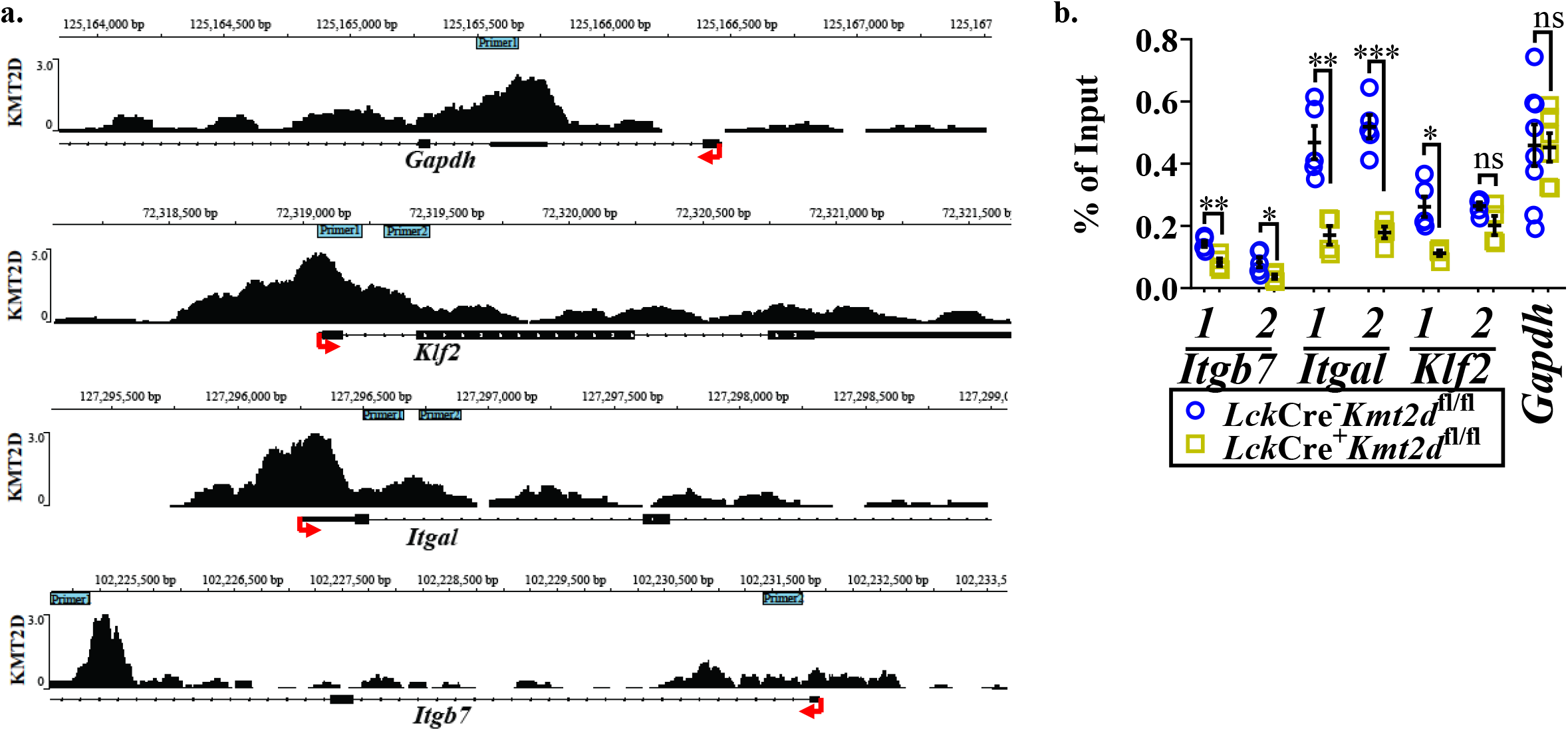
Differentially expressed egress licensing genes have KMT2D binding sites in cis-regulatory regions and demonstrate abnormal chromatin signature. (**a**) Integrative Genomics Viewer browser image of chromatin immunoprecipitation (ChIP) binding of KMT2D in control naïve CD4^+^ T cells at cis regulatory regions roughly −1 kb to + 2.5 / 5 kb around the transcription start site (TSS) region of the gene (*Itgb7, Itgal, Klf2*, and *Gapdh*) from GEO accession number: GSE69162. Each plot displays the gene location/scale and rough primer location (B; light blue/green boxes; for ChIP-PCR – **Figure-6b**) at the top and while the indicated gene, TSS indicated by red arrow, and gene structure up to the first few exons shown are on the bottom. (**b**) ChIP-PCR of H3K4me3 binding to indicated cis-regulatory regions (primer locations found in **Figure-6a**) in bulk thymocyte of egress licensing genes, *Klf2, Itgb7*, and *Itgal* as compared to that of control (*Gapdh*). The following mouse lines are displayed as cells from individual mice (n = 4 - 8) on the graphs: *Lck*-Cre^Mar^; *Kmt2d*^-SET-fl/fl^ (Cre^-^ blue circles/Cre^+^ yellow squares). Each dataset displays black lines, which represent mean ± SEM. Significance is listed on graph based on *P*-value > 0.05 (ns), *P*-value < 0.05 (*), *P*-value < 0.01 (**), and *P*-value < 0.001 (***).

To test whether abnormal chromatin marks and differential expression of *Itgal* translate into diminished protein levels, we performed flow cytometry. Our results indicate a significant decrease ITGAL expression level (decreased mean fluorescence intensity; MFI) beginning in the DP stages through SP thymocyte populations, but no major changes in ITGAL-positivity (shifts in population positive to negative ITGAL expression) in *Kmt2d*-targeted mice (**Figure 7a-c**). In contrast, both the ITGAE^+^ population and ITGAE MFI are specifically decreased in the SP thymocytes (more dramatically in CD8^+^SP, due to higher control intensity), but not DP cells (**Figure 7d-g**). Lastly, the ITGB7 MFI in both CD4^+^SP and CD8^+^SP thymocytes was decreased and with intensity decreases specifically corresponding to the CD8^+^SP ITGB7^+^ population (**Figure 7h-j**). Thus, KMT2D appears to directly control this small group of integrin signaling molecules, and these observed expression abnormalities translate to protein-level effects.

**Figure 7.**
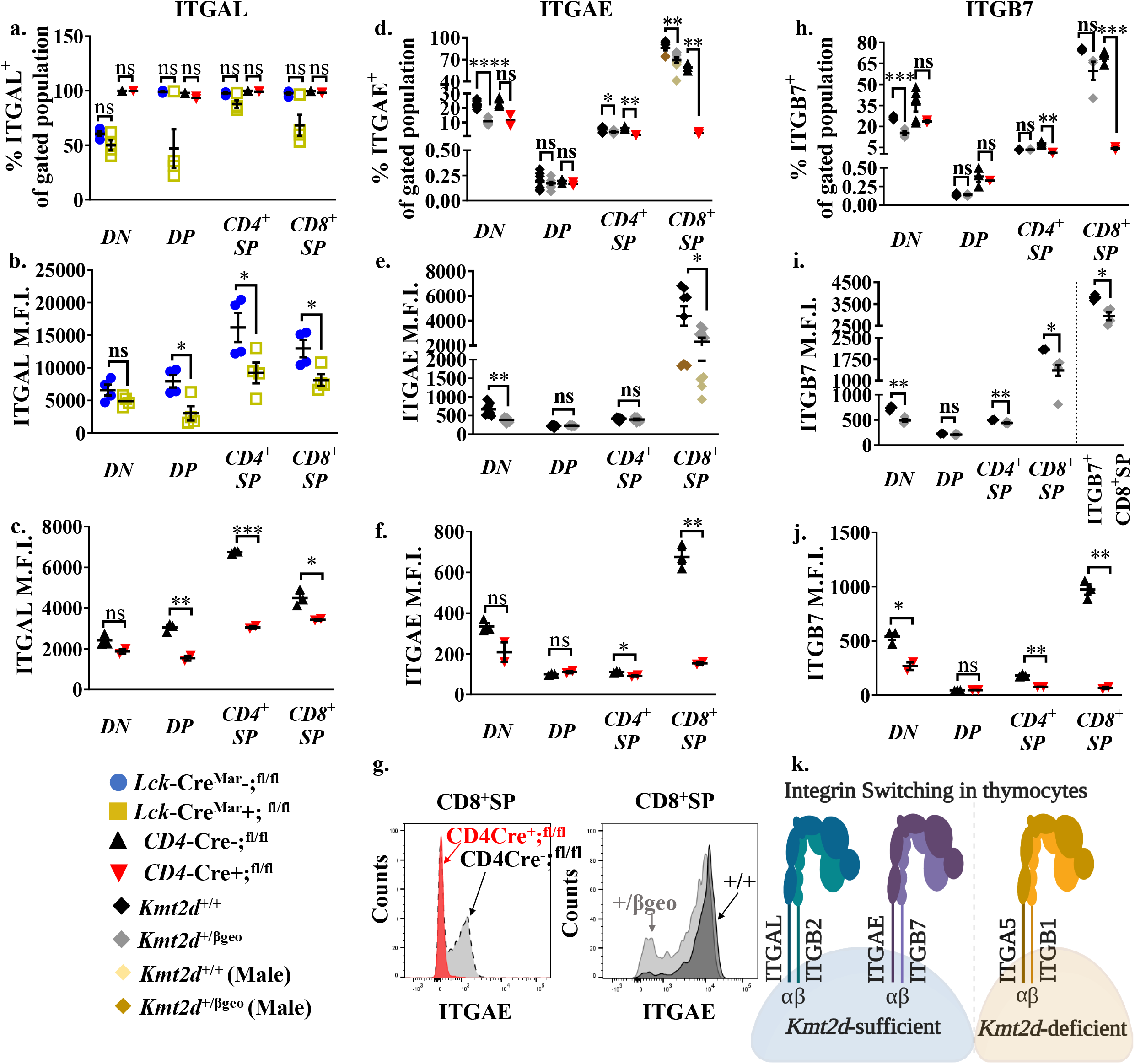
Differentially expressed integrin receptors demonstrate disrupted function. Flow cytometric population shifts across the thymocyte development. Shown as percent positive of indicated thymic population: double-negative (DN), double-positive (DP), and single-positive (CD4^+^SP, CD8^+^SP) of *ITGAL* (**a**), *ITGAE* (**d**), and *ITGB7* (**h**). Flow cytometric relative mean fluorescence intensity (MFI; i.e. protein expression level) of *ITGAL* (**b**,**c**), *ITGAE* (**e**,**f**), and *ITGB7* (**i**,**j**). MFI shown in arbitrary units of intensity (a.u.i.) for the same indicated thymic populations: DN, DP, CD4^+^SP, and CD8^+^SP. The following mouse lines (*Kmt2d^+/βgeo^* [^+/+^ black diamonds/^*+/βgeo*^ grey diamonds], *Lck*-Cre^Mar^; *Kmt2d*^-SET-fl/fl^ [Cre^-^ blue circles/Cre^+^ yellow squares] and *CD4-Cre; Kmt2d*^-SET-fl/fl^ [Cre^-^ black up-right triangles/Cre^+^ red inverted triangles]) are displayed as one marker per individual mouse (n = 2 - 9) on the graphs. Additionally, brown markers in (**d**,**e**) indicate a separately run experiment using male mice, which had lower relative ITGAE expression in wild-type compared to females; however had the same decreasing trend. Statistics displayed for all mice regardless of sex. (**g**) Representative flow histogram overlay of deficient with corresponding control littermate (y axis: Counts, as normalized to mode). Depicts of both the shift toward the negative population and decreased expression level of ITGAE in CD8^+^SP (right; black/^*+/βgeo*^ grey, left; *CD4*-Cre; *Kmt2d*^-SET-fl/fl^ with Cre^-^ black/ Cre^+^ red). (**k**) Proposed shift from ITGAL-ITGB2/ITGAE-ITGB7 toward (ITGA5)-ITGB1 in *Kmt2d*-deficient mice. Significance based on *P*-value > 0.05 (ns), *P*-value < 0.05 (*), *P*-value < 0.01 (**), *P*-value < 0.001 (***), and *P*-value < 0.0001 (****).

## DISCUSSION

KMT2D has a central role in the development of many embryonic and adult cell lineages. Herein, we identified that *Kmt2d* specifically contributes to T cell lineage development by modulating interconnected activation and maturation pathways in SP thymocytes, which ultimately alters cell fate and thymic egress. Notably, in *Kmt2d*-knockout mice, thymic M1-SP populations were increased. Moreover, both CD4^+^SP and CD8^+^SP cells had decreased expression of genes involved in activation (required for proper negative selection survival) and adhesion (most notably specific integrins, which are required for cell-cell interactions during activation and/or proper migration/localization).

Thymic compartmentalization of the medullary and cortical regions provide important signals required during particular stages of thymocyte development. Specific timing and dynamic integrin “switches” (shifts in integrin heterodimers) can have major impacts on thymocyte development by controlling thymocyte migration (adhesion, motility), activation and/or maturation capabilities. For example, ITGAE-ITGB7 binds to medullary zone epithelial cells allowing proper localization of SP thymocytes for maturation and selection ^19^. Furthermore, integrin functionality can be regulated/co-regulated by one or more combinations of: 1) integrin ligation (ITGB2 activation influences ITGB1-specific adhesion capacity and fate decisions); 2) cell activation status (resting T cells require co-expression of ITGAL-ITGB2 [LFA1]/ITGA4-ITGB1 [VLA4] for adhesion, whereas activated T cells, only require LFA1); 3) or relative proportion/combinations can control differentiation ^20^ or function (VLA4 heterodimers mediate adhesion with fibronectin in DN/CD3^lo^CD69^lo^-DP thymocytes, whereas VLA4/ITGB1-ITGA5 [VLA5] heterodimers mediate motility in mature CD3^hi^CD69^hi^ thymocytes) ^21,22^. Overall, these integrin dynamics are critical for thymocyte development.

We found that the leukocyte-specific integrins, *Itgal, Itgae*, *Itgb2*, and *Itgb7*, all demonstrate significantly reduced gene expression in T cell-driven *Kmt2d* knockout SP thymocytes. Furthermore, KMT2D regulated these integrins at cis-regulatory regions, thus its loss reduces the corresponding integrin protein expression, as we observed via flow cytometry. In contrast, we observed enhanced levels *Itgb1* expression and no significant change in one of its protein heterodimer partners, *Itga5*, whereas ITGB1’s other heterodimer partners (*Itga4/6*) decreased in CD4^+^SP thymocytes from *Kmt2d* knockouts. Together, our data suggests a potential ITGAL-ITGB2/ITGAE-ITGB7 switch toward (ITGA5)-ITGB1 in *Kmt2d* knockout SP thymocytes (**Figure 7k**).

The ITGAL-ITGB2/ITGAE-ITGB7 to (ITGA5)-ITGB1 integrin switch may have substantial implications in downstream adhesion/migration capacity, fate decision (i.e. SP maturation), and activation. One major implication could be at the immunologic synapse where LFA1/ICAM interaction occurs between the T cell and the antigen-presenting cell. Normally, the LFA1/ICAM interaction lowers the optimal T cell stimulation threshold to assist in downstream signaling of extracellular signal–regulated kinase (ERK)-1/2 during T cell activation ^23^. Reduced thymocyte LFA1 function would require a higher threshold for stimulation. A functional consequence of a pathogenic integrin switch may be insufficient or defective thymic egress. SP thymocytes may fail to egress from the thymus under a variety of specific circumstances, including; during/after negative selection secondary to altered downstream SP thymocyte TCR signaling (such as maintaining high CD69 levels) or as a results of a deficiency in egress-licensing genes normally repressed by CD69 expression (as seen in *Foxo1* or *Klf2 deficiency*) ^24–27^. Finally, KMT2D may also effect integrins indirectly via altered TCR signaling. Specifically, both RAP1A and RAP1B, which are small TCR-inducible GTPases that regulate downstream RAS-mediated activation of ERK-driven fate decisions (e.g. cell cycle, differentiation, growth), are regulated by KMT2D ^28,29^. Downstream ERK signaling is required for enhanced T cell adhesion via ITGB1 and LFA1 (mainly ITGB2) ^30^, thus creating a regulatory loop. Our data is consistent with KMT2D having a role in TCR signaling as we show that *Kmt2d*-deficient M1-SP cells accumulate within the thymus because of the blocked M1/M2-SP transition. We hypothesize this may be due to suboptimal SP thymocyte TCR stimulation impacting downstream CD69/FOXO1 signaling. This KMT2D regulation may be cell specific as *Kmt2d* deficient peripheral naïve CD4^+^ T cells (GEO: GSE69162; ^11^) do not share the similar significant integrin shifts with SP thymocytes (**Supplementary figure 11a-b**).

As multiple integrins shift together, they may be co-regulated. KMT2D thymocyte integrin co-regulation does not appear to involve genomic domain regulation as chromosomal location/loci (both murine and human) are not shared by affected integrin genes. Phylogenetically, α-integrins and β-integrins are most similar within their respective groups (**Supplementary figure 5e**). Specifically, comparative analysis of *Itgb2/Itgb7* genomic organization shows overlapping genetic structure (conservation across all 13 *Itgb7* intron-exon junctions ^31^ and high levels of amino acid homology (> 61.9%). To assess whether phylogenetically-related integrin genes may share regulatory sequences, we compared the 1-kb promoter region of the leukocyte-specific integrins and found 29.5 – 35.8% (*Itgb2/7*) and 44.6 – 49.3% (*Itgae/l*) similarity for non-identical alignments, demonstrating a potential for shared regulation. We next expanded our analysis to all putative *Kmt2d*-regulated genes identified by our analysis. Upon comparing 1-kb promoter regions upstream of the consensus transcriptional start site of all *Kmt2d*-regulated genes, we could not identify any motifs of a putative KMT2D transcriptional co-regulators from our datasets (data not shown). However, other investigators have shown a protein C-ets-1 site, retinoic acid (RA) receptor (RAR) site and a high-mobility group (HMG) binding site within both *Itgb7* and *Itga4* promoter regions ^32^. Additionally, in peripheral T cells, RA is required for *Itga4* upregulation and enhances *Itgb7* expression, but does not impact *Itgb1* ^33^. Furthermore, RAR is implicated in targeting H3K4 trimethylation to specific loci through interaction with KMT2D via the COMPASS complex ^34,35^. In total, these findings suggest that KMT2D may alter integrin expression through similar cis-regulatory elements.

Following SP integrin shifts and potential blocks at the thymic M1-SP stage in our mouse models of *Kmt2d* loss, we also observed the loss of naïve/RTE T cells in the peripheral T cell compartment and enhancement of a memory, CD44-expressing, T cell phenotype. We do not believe the enhanced peripheral memory cell phenotype in KS1 is caused by lymphopenia-driven homeostatic proliferation, as individuals with KS1 do not have significantly different percentages of total T cells; however, this could potentially be occurring in *Kmt2d* knockout models. Multiple other mouse models; however, provide useful context to our findings. *Foxo1*-deficient mice display both thymocyte egress deficiencies and decreased naïve T cell counts, indicating a potential relationship between KMT2D and FOXO1 pathways. Knockout of another histone methyltransferase, *Dot1l*, appears to phenocopy *Kmt2d* loss. Although similar in terms of the phenotypic endpoint (increased semi-mature SP thymocytes and memory-skewed peripheral T cells), loss of *Kmt2d* and *Dot1l* drive these difference by altering distinct genetic programs ^36^. Our bioinformatic analysis revealed only three shared genes are down-regulated in both knockout models (**Supplementary figure 10**). Finally, murine models deficient in *Kdm6a*, the gene haploinsufficient in individuals with KS2, has the inverse peripheral immune phenotype (T cells shift toward naïve cells with reduced memory cell counts) to *Kmt2d* loss ^37^. Beyond KMT2D, it is becoming apparent that a variety of chromatin modifiers impact lymphocyte differentiation through various non-overlapping mechanisms.

In addition to altered memory T cells, a significant increase in DNT cells was observed in both individuals with KS1 and *Kmt2d*-targeted mouse models. Of note, we observed reduced *Fasl* expression in peripheral naïve CD4^+^ *Kmt2d* knockout T cells, which suggests intrinsic loss of *Kmt2d* can decrease T cell *Fasl* expression. The individuals with KS1 within our cohort have an elevated frequency of splenomegaly compared to the general population (18.5% [3/16] compared to 10% in children and 3% adults, respectively ^38^). Additionally, we often observed splenomegaly in both constitutive haploinsufficient and hematopoietic-driven models of *Kmt2d* loss, but not in T cell-specific models suggesting T cell extrinsic *Kmt2d* loss drives splenomegaly. Increases in DNT cell counts with *Fas/Fasl* mutations (disrupted normal functioning of cellular apoptotic machinery) and elevated frequency of splenomegaly are linked to autoimmune lymphoproliferative syndrome (ALPS) (**Supplementary figure 3d/ Supplementary figure 11c**). The combination of both T cell-intrinsic and -extrinsic *Kmt2d* loss contributes to features overlapping with lymphoproliferative disorders. The splenomegaly/DNT increases are consistent with two individuals with KS investigated by Giorando *et al.;* however, in contrast to *Kmt2d*-knockout CD4^+^ T cells, no evidence of FAS deficiency was found in these individuals ^39^. Therefore, more investigation into the T cell intrinsic mechanism(s) driving DNT generation may be necessary.

In summary, we find that *Kmt2d* loss in both humans and murine KS1 models causes specific abnormalities in the T cell lineage during development. Aberrant thymocyte progression leads to downstream peripheral loss of T cells (in *Kmt2d*-knockouts), reduction of RTEs, skewing toward a memory phenotype, and increased DNT cells. *Kmt2d* has direct intrinsic effects within lymphocytes and can also exert extrinsic effects within the microenvironment. Further attention should focus on better understanding the interplay between lymphocyte *Kmt2d*-intrinsic and - extrinsic contributions (from neighboring cells, including neural crest cells, epithelial cells, B-lymphocytes, or dendritic cells) which together drive overall immune and other developmental KS1 phenotypes.

## METHODS

### Patient Consent and Sample Collection

The parents of the human subjects provided written consent for enrollment in this study (Cincinnati Children’s Hospital Medical Center [CCHMC] Institutional Review Board [IRB], protocol #2012-4636 and/or protocol #2020-0570). All KS1 cohort peripheral blood lymphocyte subpopulation analysis was performed in the CCHMC Diagnostic Immune Laboratory and the collected data were compiled from review of medical records. All clinical evaluations with the exception of DNT cells were performed at Clinical Laboratory Improvement Amendments (CLIA)–certified clinical laboratories and reported with the age-matched normal ranges. DNT cell gating is not a standard lymphocyte cell population assessed during routine clinical lymphocyte analysis, but rather a specialized cell type assessed only under particular situations. DNT populations have been described across all age ranges to be variant above 2.5% or 1.5% within CD3^+^ populations or with all lymphocyte gating strategies, respectively ^40,41^. All 16 KS1 cohort patients underwent *KMT2D* sequencing (reference transcript: NM_003482). KMT2D domains were mapped according to UniProt Database (O14686). Protein mutations and type (frameshift, nonsense, missense, splice variants) associated to the individual’s recorded DNA mutation were generated via VarSome (https://varsome.com/) ^42^ for the Lolliplot (https://www.cbioportal.org/mutation_mapper) ^43,44^. Lolliplots were manually altered in order to include additional variant information through color and shape. Values found within (closed) and outside (open) the reference range are both displayed on the plots. The significance of the individuals falling within the reference range was determined by the exact binomial test (using *P*-value = 0.975, as the reference range displayed is 2.5 - 97.5 % confidence) and the *P*-value generated is listed on the graphs in **Table 1**. The lower limit of normal (LLN) for assessment of cardiac surgery is calculated using the KS1 value divided by the lowest value (2.5 confidence value) of the reference range.

### Mice

Mice were housed in accordance with NIH guidelines, and experimental protocols were approved by the Institutional Animal Care and Use Committee of CCHMC and/or Johns Hopkins University. The thymus, spleen, and/or blood were collected for flow cytometry and phenotyping analysis. Magnetic bead sorting was performed to sort out SP cells or peripheral T cell subsets for assessment of Cre recombinase efficiency and for RNA isolation, RNA-seq library preparation, and sequencing. *Kmt2d^+/βgeo^* (also known as *Mll2Gt^(RRt024)Byg^*) mice (genetrap) were obtained from Bay Genomics (Berkeley, Calif) and contain a powerful splice enhancer integrated in between exons prior to the SET domain. These mice show many features in common with patients with KS ^45^. *Kmt2d^+/βgeo^* animals were backcrossed onto a pure C57BL/6 background (verified by SNP array via Taconic Labs) and are maintained through breeding heterozygous carriers with fresh C57BL/6J (B6; Jackson Laboratories) as homozygous *Kmt2d ^βgeo/βgeo^* mice are embryonic lethal. *Kmt2d*^-SET-fl/fl^, (also known as *Mll4*^-SET-fl/fl^ mice) a kind gift from Dr. Kai Ge (NIH) have been previously described ^46,47^, were crossed with Cre recombinase (Cre) under various promoters. This mouse strain is not the same as the publicly available *Kmt2d* ^taL.1Kaig/J^ (also known as *Mll4*^fl/fl^) mouse from Jackson laboratories (Strain #032152), which we refer to in this document as *Kmt2d* ^exon16-19-fl/fl^ to differentiate it from *Kmt2d*^-SET-fl/fl^. Briefly, these mice harbor LoxP sites around *Kmt2d* exons 50 and 51 and disrupt the SET domain through a frameshift causing a stop codon at exon 52 upon exposure to Cre, which leads to a truncated protein (276 amino acid truncation at the carboxy-terminal end).

Conditional mice will sustain *Kmt2d* loss throughout thymocyte development and into the mature T lymphocytes in the periphery, thus allowing us to determine the role of *Kmt2d* in T cell development. *Vav1*-iCre (a kind gift from Senad Dianovic, CCHMC, but can also be purchased from The Jackson Laboratory, Bar Harbor, USA; Strain #008610) turns on at the hematopoietic stem cell stage, whereas the two additional models to conditionally delete T cells, *Lck*-Cre^Mar^ (The Jackson laboratory; Stain #003802) reportedly turns on around DN3 ^12^ and *CD4*-Cre (a kind gift from David Hildeman, CCHMC but can also be purchased from The Jackson laboratory; Strain #022071) turns on around DN4 (^12^; see **Supplementary Figure 2**). In one experiment, *Vav1*-iCre; *Kmt2d*^-SET-fl/fl^ mice were sensitized with 100 μg ovalbumin (OVA #A5503; Sigma, St. Louis, USA) and 1 μg cholera toxin (CT #101B; List Biological Laboratories Inc., Campbell, USA) in 50 μl sterile Dulbecco’s Phosphate Buffered Saline without either Calcium or Magnesium Chloride (DPBS, #14190-144; ThermoFisher Scientific, Waltham, USA) intratracheally once a week for three weeks straight, waited two weeks then spleens were harvested. Spleen weights and size by photograph were recorded.

All mice used in these studies were approximately 1 - 6 months old. Mice were sacrificed via intraperitoneal injection of sodium pentabarbitol (65 mg mL^-1^; 200 – 300 uL) after which the thymus, spleen, mesenteric lymph node and/or blood were collected. Using a 3-mL syringe plunger (#309656; Becton Dickinson [BD], Franklin Lakes, USA), the organs were mashed through a T-cup strainer (#352350; Corning Inc., Glendale, USA) using 5 – 10 mL of 0.5% DNase I (DN25-1G; Sigma, St. Louis, USA) in complete RPMI 1640 (cRPMI; RPMI 1640 [#10-040-CV; Corning/Mediatech Inc., Manassas, USA] containing 10% heat inactivated Fetal Bovine Serum [#16140-071; Invitrogen]). Peripheral blood was collected from live mice for analysis via submandibular phlebotomy into K2EDTA tubes (Microtainer, #365974; BD). Cells were collected and spun down at 500 *g* x 5 minutes. The 1 million cells collected from tissues or blood were strained and used for flow cytometry analyses or for recent thymic emigrant T-cell receptor excision circles assessment.

### Flow Cytometry Analysis

Cells were spun down at 500 g for 5 minutes to pellet and resuspended in 5 mL of cRPMI. Cells were stained using a dilution of 1:400 of anti-mouse CD16/CD32 (Fc Shield, #70-0161-U100; TONBO biosciences, San Diego, USA) in cRPMI for 15 minutes, washed with cRPMI, spun down, and incubated with the following antibodies in cRPMI at 1:200 for 30 minutes: CD4-BV650 (#100469; Biolegend, San Diego, USA), CD8-BV605 (#100744; Biolegend), CD25-PeCy7 (#102016; Biolegend), CD44-AF488 (#103016; Biolegend) and additional panel antibodies (TCRβ, ITGAL, ITGAE, MHCI, CD69, CCR7, CD24) or peripheral (splenic) antibody panel; see **Supplementary table 1** and **Supplementary figure 12** for antibody information and gating strategies, respectively. For analysis only: cells were washed with 1X DPBS (10X DPBS diluted with Milipore water 1:10; #14200-075; ThermoFisher Scientific) twice then stained at 1:1000 with Zombie UV Fixable Live/Dead (blue-fluoresent reactive dye #L23105A; Thermofisher Scientific) for 15 minutes at room temperature. Cells were washed, spun down and fixed with either 4% diluted paraformaldehyde (#15710-S; Electron Microscopy Sciences, Hatfield, USA) or with Cytofix/Cytoperm (#554722; BD) for 20 – 30 minutes, washed with 1X DPBS, spun down, and stored in 1X DPBS or cRPMI until run on the LSR Fortessa I flow cytometer (BD, maintained/run at CCHMC) or FACSverse (BD, run at JHU). Flow cytometry files were analyzed using FlowJo (Treestar LLC, now BD, Ashland USA).

### Magnetic Bead Sorting

Cells were then spun down at 500 x *g* for 5 minutes to pellet and resuspended in 5 mL of cRPMI. Single cells were depleted from both the CD4^+^ double positive (DP) and CD4^+^ single positive (SP) cells using a CD4^+^ (L3T4) positive selection kit (Microbeads; #130-117-043; Miltenyi Biotec, Gaithersburg, USA) flow-through (this contains both double-negative [DN] and CD8^+^SP cells). CD8^+^ SP thymocytes were isolated using the CD8a^+^ (Ly-2) positive selection kit per manufacturer’s instructions (Microbeads, #130-117-044; Miltenyi Biotec). Flow-through resulted in the DN population, whereas the magnetic elute resulted in the CD8^+^SP population. Conversely, CD4^+^SP isolation was used in the opposite microbead sequence (CD8 then CD4). Briefly, both these kits used a 10 minute stain with either CD4^+^- or CD8^+^-specific microbeads followed by applying cells to the magnet-associated column, thereby capturing the CD4^+^ or CD8^+^ cells, relative to the antibody-microbead used.

### Recombination Assessment of Cre

Recombination of magnetic sorted thymocyte and peripheral cells (pool from the spleen and lymph-nodes) CD4^+^ and CD8^+^ cells (see **Supplementary figure 2A/B** for Cre recombination activity documented in prior literature [**Supplementary figure 2A**] and location of our designed primers and predicted outcomes [**Supplementary figure 2B**/ **Supplementary table 2**]) was assessed by determining the expression or loss of exon 50/SET domain through genomic DNA quantitative polymerase chain reaction (PCR). CD4^+^ and CD8^+^ cells were sorted through magnetic bead separation (see Magnetic Bead Sorting procedure above) and genomic DNA was isolated via QIAamp DNA micro kit (#56304; Qiagen LLC, Germantown, USA) per manufacturers’ directions and quantified optically by Qubit 2.0 Fluorometer (#Q32866; ThermoFisher Scientific) using Qubit dsDNA HS Reagent/Buffer (#Q32854A/Q32854; ThermoFisher Scientific). A quantitative PCR reaction was performed using 10 ng (in 3 μL volume) of extracted DNA per reaction, 2 μL of combined 1μM forward and reverse primers (see **Supplementary table 2**) and 5 μL SYBR green master mix (#A25742; Applied biosystems, Waltham, USA) and run on the QuantStudio7Flex (Applied Biosystems) or ABI ViiA7 (Applied Biosystems) in a MicroAmp Optical 384-well reaction plate (#A4309849; Applied biosystems). The CT values were then placed in the following equation: 2^-(Mutant[CT of *Kmt2d*-Exon50 / CT of*Kmt2d*-Total R20] – Control[CT of *Kmt2d*-Exon50 / CT of *Kmt2d*-Total R20])^ for recombination assessment.

### T cell receptor excision circles (TRECs) Analysis of Recent Thymic Emigrants (RTE)

Within developing T cells, rearrangement of the gene segments encoding the T-cell receptor (TCR) occurs, resulting in chromosomal sequences that are excised to produce episomal DNA by-products, called TCR rearrangement excision circles (TRECs). TRECs are stable, not duplicated during mitosis, and diluted out with each cellular division; therefore, TRECs serve as markers for recent thymic emigrants (RTEs) in the peripheral blood. To detect RTE, TREC numbers are analyzed from peripheral blood and normalized to the T cell numbers from the blood via flow cytometry gating. Briefly, peripheral whole blood was collected in K2EDTA Microtainer tubes (BD; #365974) or with 50 – 100 μL of Sodium Heparin–coated tubes (#540-15; Fresenius Kabi, Lake Zurich, USA), subsequently stained with CD45-PE-cy7 (#368531; BioLegend) or CD45-Alexa700 (#103113; Biolegend) and CD3-FITC (#130-080401; Miltenyi Biotec), fixed/lysed with Cytofix/Cytoperm (554722; BD), and stored in 1X PBS until run on the flow cytometer (BD LSR Fortessa/BD FACSverse). Gating percentage was used to normalize the data to CD45^+^ CD3^+^ total T cells. DNA from 100 μL of whole blood was extracted using a QIAamp DNA micro kit per manufacturers’ directions and quantified optically by Qubit 2.0 Fluorometer, as done in “**Recombination Assessment of Cre**” section. A PCR reaction was performed using 12.5 – 40 ng of extracted DNA per reaction and PowerUp SYBR green master mix (#A25742; Applied biosystems). Mouse primer Ca was used as an internal control (Mouse Ca Forward: 5’-TGACTCCCAAATCAATGTG-3’; Mouse Ca Reverse: 5’-GCAGGTGAAGCTTGTCTG-3’), whereas mouse primers for TREC were used to quantify TREC (Mouse sjTREC Forward: 5’-CCAAGCTGACGGCAGGTTT-3’; Mouse sjTREC Reverse: 5’-AGCATGGCAAGCAGCACC-3’). Cycling was as follows: 2 minutes at 50°C, 2 minutes at 95°C, 40 cycles of: 15 seconds at 95°C and 1 minute at 60°C. After the CTs were extracted, the differences in CT (delta CT) from Ca and sjTREC were calculated and run through the equation sjTREC / percent CD45^+^CD3^+^ cells to normalize to the number of T cells.

### RNA Isolation, Library Preparation, and Sequencing

RNA from sorted *Kmt2d*-knockout and littermate control thymic cells was isolated by centrifugation to pellet and resuspended in 300 μL of Protection Buffer (Monarch RNA Isolation Kit, #T2010S; New England Biolabs Inc. (NEB), Ipswich, USA;) and stored at −80°C until isolation. RNA was subsequently isolated following the manufacturer’s instructions of the Monarch RNA Isolation Kit (#T2010S; NEB). RIN scores and concentration were performed using High Sensitivity RNA ScreenTape (#5067-5579/#5190-6507; Agilent Technologies, Santa Clara, USA) and run using the Agilent Technology software with an electronic ladder. Quality RIN scores above 7 were used for RNA-Seq library preparation. Samples were stored at −80°C until library construction. Library construction (input: 40 ng per CD8^+^SP sample) was performed using a NEBNext Poly(A) Magnetic Isolation Module (#E7490; NEB) followed by a NEBNext Ultra II Directional RNA library prep kit for Illumina (#E7760 and/or #E7770 with #E7765; NEB) with size selection by AMPure XP (#Ab3881; Beckman Coulter, Brea, USA) beads, according to manufacturers’ protocols. Library quantification and quality checks were done using KAPA Library Quantification Kit for Illumina (#KK4824; Kapa Biosystems, Cape Town, South Africa), High Sensitivity D1000 DNA Kit on BioAnalyzer (#5067-4626/#5190-6502; Agilent Technologies). Paired-end 50-bp read sequencing was obtained for pooled libraries using Novaseq 6000 (Illumina, San Diego, USA).

#### RNA-Seq Analysis

Transcriptomic data collected by RNA-Seq was analyzed to determine the genes that were present in each sample, their expression levels, and the differences in expression levels between *Kmt2d*^SET-fl/fl^ *Cre^+^* conditional mutant and *Kmt2d*^SET-fl/fl^ control (CD4^+^SP *Lck*-Cre^Mar^, CD8^+^SP *Lck*-Cre^Mar^, CD8^+^SP *Vav1*-iCre, and also utilized publically accessible data CD8^+^SP from *Lck*-Cre *Dot1l*^*f*l/fl^ GEO: GSE138910, and CD4^+^ peripheral naïve T cells from *CD4*-Cre *Kmt2d*^exon16-19-fl/fl^ GEO: GSE69162). RNA-Seq analysis was performed using three methods. The results of the analysis from the first analysis program, SciDAP (https://scidap.com/), are used for the figures; the two alternative methods (Salmon [2] and HISAT2 [3] alignment tools) were used for verification (data not shown).

RNA-Seq Analysis Alternative (for confirmation): For Salmon (2) and HISAT2 (3) alignment tools: following quality checking with the software Fastqc, reads were trimmed with trim-galore/0.5.0 to remove reads with adaptors.

(2) Salmon alignment tool. We built a mouse index using FASTA file of all mouse cDNA sequences downloaded from Ensembl (ftp://ftp.ensembl.org/pub/release-91/fasta/mus_musculus/cdna/Mus_musculus.GRCm38.cdna.all.fa.gz) and pseudo-mapped the RNA-Seq reads with Salmon (v1.1). Subsequently, resulting transcript quantifications were imported into R to get gene-level counts, using the tximport R package. The differential analysis was performed with DESeq2, retaining counts greater than 10.

(3) HISAT2 alignment tool. We build a mouse index using FASTA file downloaded from Ensembl (ftp://ftp.ensembl.org/pub/release-101/gtf/mus_musculus/Mus_musculus.GRCm38.101.gtf.gz) and mapped the reads with the alignment tool HiSat2/2.1.0. The generated Sam files were processed using samtools/1.9. The reads mapped to feature (exon) and meta-feature (gene) were counted with the feature Counts function from Subread/1.6.3 using an annotation file downloaded from Ensembl (ftp://ftp.ensembl.org/pub/release-101/gtf/mus_musculus/Mus_musculus.GRCm38.101.gtf.gz). This count matrix served as the input for the differential analysis, which we performed using DESeq2. For (2) and (3): Differentially expressed genes were further analyzed for the enriched pathways with an online tool, WEB-based Bene SeTAnaLysis Toolkit (webgestalt.org).

Before uploading into SciDAP, catenation of the two files from each lane were merged into the same file (R1 and R2 merged; L1 and L2 merged). Within the bioinformatics website SciDAP: we imported the merged FASTQ files, trimmed adapters from the files (https://github.com/datirium/workflows/workflows/trim-rnaseq-pe.cwl) and aligned samples to the mouse genome (Mus Musculus [mm10] genome reference consortium mouse build 38, strain: C57BL/6J. 2012/01/09) and generated genome indices using STAR v2.5.3a (03/17/2017) ^48^, as previously described ^49,50^. The resulting sample files created from the alignment were then combined to perform differential gene expression analysis based on the negative binomial distribution (DESeq2) analysis pipeline (https://github.com/Barski-lab/workflows-datirium/workflows/deseq.cwl). Each set of data (*Kmt2d*^SET-fl/fl^ *Cre^+^*) with their corresponding littermate controls (*Kmt2d*^SET-fl/fl^) were used in their specific DESeq2 runs (i.e. 1. CD4^+^SP *Lck*-Cre^Mar^, 2. CD8^+^SP *Lck*-Cre^Mar^, 3. CD8^+^SP *Vav1*-iCre). Additionally, publically available datasets were compared with our DESeq2 datasets. DESeq2 was performed on the naïve *Kmt2d*-knockout CD4^+^ T cells with their experimental controls (GEO accession: GSE69162; **Supplementary figure 5/9**) and *Dot1l*-knockouts with their experimental controls (GEO accession: GSE138910; **Supplementary figure 10**). Data files were exported from SciDAP and filtered within Microsoft Excel using the following criteria: RPKM ≥ 5 in at least one condition, adjusted *P*-value ≤ 0.01, and Log2 fold change ≥ 0.59 or ≤ −0.59 to be considered “up-regulated” or “down-regulated,” respectively, in wild-type compared to *Kmt2d*-deficient/mutant mice. Gene visualization (up-regulated/down-regulated vs. control littermates) is displayed via Volcano plot (VolcaNoseR; https://goedhart.shinyapps.io/VolcaNoseR/; ^51^)

### Visualization of Down-regulated Genes in *Kmt2d*-knockout Compared to Littermate Control Gene Sets

The resulting down-regulated lists (down-regulated in *Kmt2d*-deficient compared to control mice) were placed into a Venn diagram (Venny v2.1; https://bioinfogp.cnb.csic.es/tools/venny/index.html; ^52^) to determine the overlapping up-regulated genes in the *Kmt2d*^SET-fl/fl^ controls from all three thymocyte datasets (1. CD4^+^SP *Lck*-Cre^Mar^, 2. CD8^+^SP *Lck*-Cre^Mar^, 3. CD8^+^SP *Vav1*-iCre). The resulting genes were copied into ToppGene Suite (https://toppgene.cchmc.org/; ^53–56^) using the program ToppFun and the manufacturer’s recommended settings and analyzed for GO terms (with cut-offs being false-discovery rate Benjamini and Yekutieli [FDR B&Y] = 0.05). The heat map was generated via importing a table of the GO select gene list generated from combined biological processes from the top hits that were found in both adhesions and activation categories with their corresponding DESeq2 average values within Microsoft Excel into Morpheus (https://software.broadinstitute.org/morpheus).

All individual sample RPKM values were run using the Feature expression merge pipeline, a program that combines RPKM expression from several experiments, from SciDAP. RPKM were graphed using PRISM (Graphpad), with *P*-values listed from DESeq2-generated adjusted *P*-values.

### Proposed Transcription Factor Binding Promoter Region of Down-regulated Genes

To understand potential transcription factors binding within the promoter region of down-regulated genes in the following variants: 1. CD4^+^SP *Lck*-Cre^Mar^, 2. CD8^+^SP *Lck*-Cre^Mar^, and 3. CD8^+^SP *Vav1*-iCre compared to their littermate controls (via DESeq2 criteria), we used the defined upstream 1-kb from the transcriptional start site (promoter region) locations from our filtered down-regulation in variant gene lists and within SciDap using the Motif Finding with HOMER with random background regions ^57^. The two resulting transcription factor binding motif lists (known and *de novo*) from each dataset were generated. The proposed transcription factors were filtered against our DESeq2 RPKM values for expression. These transcription factors were compared between known and *de novo* for each dataset for validation, and compared across datasets. To determine if a transcription factor binding motif is true (not an artifact), the motif is required to be consistently enriched across the models and contain the length/complexity needed to accurately assign.

### Chromatin immunoprecipitation (ChIP) –Polymerase Chain Reaction (PCR)

Thymi were harvested and single cell suspensions were created as described above. As previously described by Sailaja *et al*. ^58^, 8 x 10^7^ total thymocytes were cross-linked with 1% formaldehyde in PBS with 0.5% Bovine Serum Albumin at room temperature for 10 minutes, followed by the addition of glycine in water solution to a final concentration of 125 mM glycine for 5 minutes. After a two washes with ice-cold PBS and their subsequent centrifugations at 300 *g* for 7 minutes each, the cells were lysed with ChIP-Lysis buffer (10 mM EDTA, 1% [w/v] SDS, 50mM Tris-HCl, pH 7.5, 1 tablet Protease Inhibitor Cocktail [#11836170001; Roche, Mannheim, Germany] and 0.1 mM phenylmethylsulfonyl fluoride [PMSF], followed by sonication in a Biorupter water bath of 5 cycles of 5 minutes, set on HIGH, 30 s ON, and 30 s OFF. Twenty-five ug of DNA diluted in ChIP dilution buffer (0.01% [w/v] SDS, 1.1% [v/v] Triton, 1.2 mM EDTA, 16.7 mM Tris-HCl, pH 8.1, 167 mM NaCl) were precipitated overnight at 4°C with H3K4me3 antibody (#9727; Cell signaling technology, Danvers, USA) or normal rabbit IgG (considered “input,” # 2729; Cell signaling technology). PCR of the precipitated product (5 ng) was performed using a reaction with 5 μL SyBR green (#A25742; ThermoFisher Scientific) and 0.5 μL of 10 μM primer listed in the **Supplementary table 3**. Primer sequences were designed based off the 8 week Thymus H3K4me3 Histone Modification by Chip-seq Signal from ENCODE/LICR located under the track name Thymus H3K4me3 in the Mouse July2007 (NCBI37/mm9) assembly on UCSC genome browser ^59,60^. Primers were designed with the intention that the amplicons would span the H3K4me3 binding peak.

### Chromatin immunoprecipitation (ChIP)

IVG genome browser (accessed via SciDAP Trim Galore ChIP Seq Pipeline Single-Read; mm10) was used to locate KMT2D binding in cis-regulatory regions of the egress licensing genes within −1-kb of transcriptional start site (TSS) into the gene body first few exons (up to 5-kb) in naïve CD4^+^ T cells from a publically accessed ChIP-Seq file SRR2037221 analyzed using corresponding input file SRR2037222 ^61,62^. In silico PCR, UCSC genome browser with mm10 genome, was used to generate the primer location (added to the image in blue) from the ChIP-PCR primer sequence ^59^.

### Assessment of Gene and Protein Sequence Similarities

Murine ITGAE (Gene ID: 16407; Protein Genbank ID: ABD49099.1), ITGAL (16408; AAI45803.1), ITGB2 (16414; AAI45645.1), ITGB7 (16421; EDL03997.1) protein and gene sequences were obtained from the NCBI database for genetic and protein similarities. For comparison of the cis regulatory region (1001 base pairs upstream from the transcriptional start site sequence was determined using UCSC genome browser with mm9 alignment and the DNA sequence generated. and run through LALIGN Pairwise Sequence Alignment (https://www.ebi.ac.uk/Tools/psa/lalign/; a program that finds internal duplications by calculating non-intersecting local alignments of protein or nucleotide sequences ^63^).

### Data accessibility

The RNA-seq data discussed in this publication have been deposited in NCBI’s Gene Expression Omnibus (GEO) ^64^ and are accessible through GEO Series (GSE) accession GSE205285 (https://www.ncbi.nlm.nih.gov/geo/query/acc.cgi?acc=GSE205285). *GSE205285 access will be provided upon acceptance of this manuscript or upon reviewer request*. Additionally, the publically available datasets from GSE accessions: GSE69162 and GSE138910 were included in our analyses and are linked to GSE205285. All data that support the finding(s) of this study are available from the corresponding authors upon reasonable request.

## ACKNOWLEDGMENTS and CONFLICT OF INTEREST

Special thanks goes to Casey Wells (CCHMC) for assistance with flow cytometry files and Shawna Hottinger (CCHMC) for editorial assistance. We thank David Hildeman (CCHMC) for fruitful discussions throughout this project. AB is a co-founder of Datirium, the developer of Scientific Data Analysis Platform (https://SciDAP.com) used here for processing RNA-Seq data. AWL is an employee of Amgen Inc. HTB is a consultant for Mahzi therapeutics. The authors have no additional competing potential financial of interests. HTB and LZ are funded by the Louma G. Foundation, which specifically supported this work. HTB is also funded by The Icelandic Research Fund (#217988, #195835, and #206806) and the Icelandic Technology Development Fund (#2010588-0611). The work of SJP, AWL, and AB was funded by a Center for Pediatric Genomics (CpG) grant from the Cincinnati Children’s Research Foundation and supported by the Center for Stem Cell and Organoid Medicine (CuSTOM) at Cincinnati Children’s Hospital Medical Center. Flow cytometric data generated by SJP were acquired using equipment maintained by the Research Flow Cytometry Core in the Division of Rheumatology at Cincinnati Children’s Hospital Medical Center (CCHMC).

---

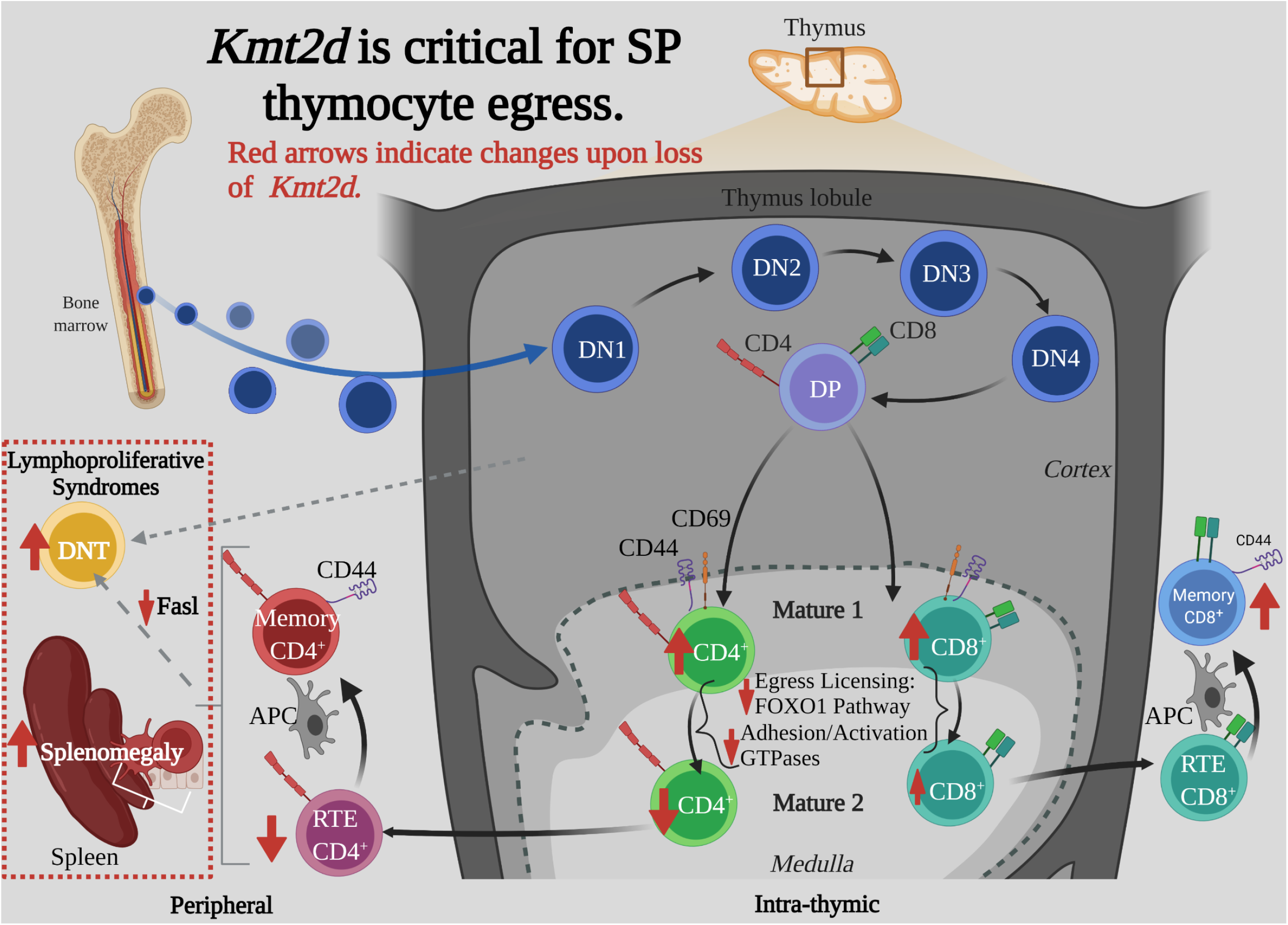

