## Supplementary Figures for "KMT2D Regulates thymic Egress by Modulating Maturation and integrin Expression"

**Supplementary table 1: Flow cytometry panels**

| Antibody | Fluorophore | Company | Catalogue # | Clone | Panel |
|---|---|---|---|---|---|
| CD3e | PE-Cy5 | Biolegend | 100310 | 145-2C11 | † Peripheral |
| CD4 | eFluor650 | Biolegend | 100469 | GK1.5 | † Peripheral |
| CD8 | eFluor605 | Biolegend | 100744 | 53-6.7 | † Peripheral |
| CD44 | AF488 | Biolegend | 103016 | IM7 | † Peripheral |
| CD62L | APC-Cy7 | Biolegend | 104428 | MEL14 | † Peripheral |
| Live/Dead (Blue-fluorescent reactive dye) | ZombieUV | Invitrogen/ Thermofisher Sci. | L23105A | n/a | † Peripheral/ ‡ §Semimature/Integrin/ ¶ Thymus Population |
| CD69 | PE | Biolegend | 104507 | H1.2F3 | ‡ Semimature |
| CCR7 | PerCP (Cy5.5) | Biolegend | 120115 | 4B12 | ‡ Semimature |
| MHCI | APC | eBioscience/Invitrogen/ Thermofisher Sci. | 17-5958-82 | AF6-88.5.5.3 | ‡ Semimature |
| TCRbeta | FITC | Biolegend | 109205 | H57-597 | ‡ Semimature |
| CD4 | PE-Cy7 | BD Bioscience | 561099 | Rm4-5 | ‡ Semimature |
| CD8 | V450 | BD Bioscience | 560471 | 53-6.7 | ‡ Semimature |
| CD4 | BV650 | Biolegend | 100469 | GK1.5 | §Semimature/Integrin |
| CD8 | BV605 | Biolegend | 100744 | 53-6.7 | §Semimature/Integrin |
| CD24 | Pacific Blue | Biolegend | 101820 | M1/69 | §Semimature/Integrin |
| TCRbeta | BUV737 | Biolegend | 612821 | H57-597 | §Semimature/Integrin |
| ITGAL | APC | Biolegend | 101120 | M17/4 | §Semimature/Integrin |
| ITGAE | PerCP-eF710 | eBioscience/Invitrogen/ Thermofisher Sci. | 46-1031-82 | 2E7 | §Semimature/Integrin |
| ITGB7 | APC | Biolegend | 321207 | FIB504 | §Semimature/Integrin |
| CD4 | BV650 | Biolegend | 100469 | GK1.5 | ¶ Thymus Population |
| CD8 | BV605 | Biolegend | 100744 | 53-6.7 | ¶ Thymus Population |
| CD25 | PE-Cy7 | Biolegend | 102016 | PC61 | ¶ Thymus Population |
| CD44 | AF488 | Biolegend | 103016 | IM7 | ¶ Thymus Population |
| TCRbeta | BUV737 | Biolegend | 612821 | H57-597 | ¶ Thymus Population |

† Peripheral : *Kmt2d*$^{+/\beta geo}$ and *CD4*-Cre; *Kmt2d*$^{SET\text{-}fl/fl}$ mouse lines.
‡ Semimature : *Lck*-Cre$^{Mar}$; *Kmt2d*$^{SET\text{-}fl/fl}$ mouse line.
§ Semimature/Integrin : *Kmt2d*$^{+/\beta geo}$ and *CD4*-Cre; *Kmt2d*$^{SET\text{-}fl/fl}$ mouse lines.
¶ Thymus Population : *Kmt2d*$^{+/\beta geo}$ and *CD4*-Cre/Vav1-iCre; *Kmt2d*$^{SET\text{-}fl/fl}$ mouse lines.

**Supplementary table 2: q-PCR Recombination Primers**

| Primer Forward/ Reverse Name | Primer Sequence | Expressed |
|---|---|---|
| *Kmt2d*-Exon50 -For | 5'-CTG TGT GGA ACC GCA TCA TTG -3' | Control Expressed<br>KO Not Expressed |
| *Kmt2d*-Exon50 -Rev | 5'-CAG CCC AAA GAG TTC TTC GCC -3' | |
| *Kmt2d*-Total R2 -For | 5'-TCT CCC GAG ACT CAG TCA CTG-3' | Control Expressed<br>KO Expressed |
| *Kmt2d*-Total R2 -Rev | 5'-CAG TTG AGC TAG TCA AGT GAT T -3' | |
| *Kmt2d*-P3e-For | 5'-TTC CAT AGC CAT TGC TCA AA -3' | Control Not Expressed<br>KO Expressed |
| *Kmt2d*-F1r-Rev | 5'-GAA CGG ATC CAA GCT TAT GC -3' | |
| *Kmt2d*-F1-For | 5'-GCA TAA GCT TGG ATC CGT TC -3' | Control Expressed<br>KO Expressed |
| *Kmt2d*-R2-Rev | 5'-CTG AAG TTT GGG AGG GTC AC -3' | |

**Supplementary table 3: ChIP-PCR Primers**

| Primer | Strand | Sequence | Location of Seq in mm10 (UCSC In-Silico PCR) |
|---|---|---|---|
| ***Klf2* Set 1** | *Sense* | GCCTATCTTGCCGTCCTTT | chr8:72319092+72319237 146bp |
| | *Anti-Sense* | TGGACCTTGTCATCTCCAGTA | |
| ***Klf2* Set 2** | *Sense* | CTTGAGGGCCTAGTTGTTAGAC | chr8:72319344+72319453 110bp |
| | *Anti-Sense* | CCGCCTCGGGTTCATTT | |
| ***Itgb7* Set 1** | *Sense* | CTCAGGGTCTTCAAGGTTACAG | chr15:102225073+102225193 121bp |
| | *Anti-Sense* | GTCGTCTTTATCGTCCTCCTTC | |
| ***Itgb7* Set 2** | *Sense* | CTGGGAGAAGCACAAGAGTAAG | chr15:102231561+102231678 118bp |
| | *Anti-Sense* | CTCCTGCTGCTCATCCATTT | |
| ***ItgaL* Set 1** | *Sense* | TCCAGCTGTTTGGTAGGAAATC | chr7:127296512+127296615 104bp |
| | *Anti-Sense* | GAGTGGCTCCTCATCTTCTTTG | |
| ***ItgaL* Set 2** | *Sense* | AGTGAGCCTTCACGTGTTTAG | chr7:127296735+127296819 85bp |
| | *Anti-Sense* | CAAAGGAGGTGGAGTAGAAACC | |
| ***Gapdh*** | *Sense* | CTC CTG CGG CCC ACT CCG CGA | chr6:125165524+125165626 103bp |
| | *Anti-Sense* | CGG ACTGCA GCC CTC CCT GGT | |

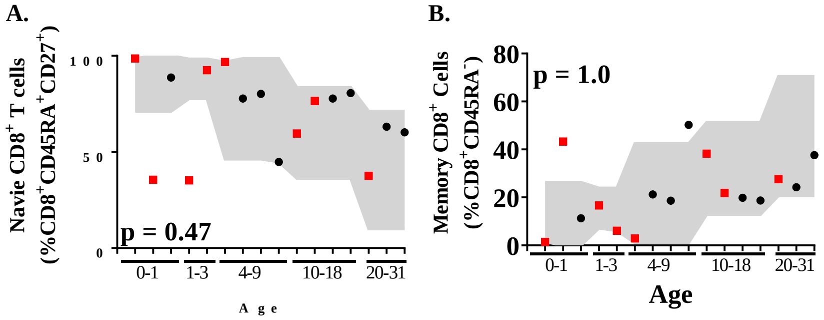


**Supplementary figure 1. Naïve and memory CD8$^{+}$ T cells in individuals with KS1.** (**a**) Data from clinical flow cytometry. Naive CD8$^{+}$ T cells (CD8$^{+}$CD45RA$^{+}$CD27$^{+}$) as a percent of CD8$^{+}$ cells. (**b**) Data from clinical flow cytometry. Memory CD8$^{+}$ T cells (CD8$^{+}$CD45RA$^{-}$) as a percent of CD8$^{+}$ cells. Values found within grey region (closed marker; represents reference population range [2.5% - 97.5%] for the healthy age-matched individuals). Significance was determined using the binomial test, with the null hypothesis being that individuals with KS1 have a same probability of falling outside the reference range equal to 5%, that is, the same as healthy age-matched individuals. The resulting *P*-values were subsequently adjusted for multiple testing with the Bonferroni method. Cardiac surgery (red square; non-surgery: black circle), required by many KS1 individuals, can potentially remove and/or damage the thymus.

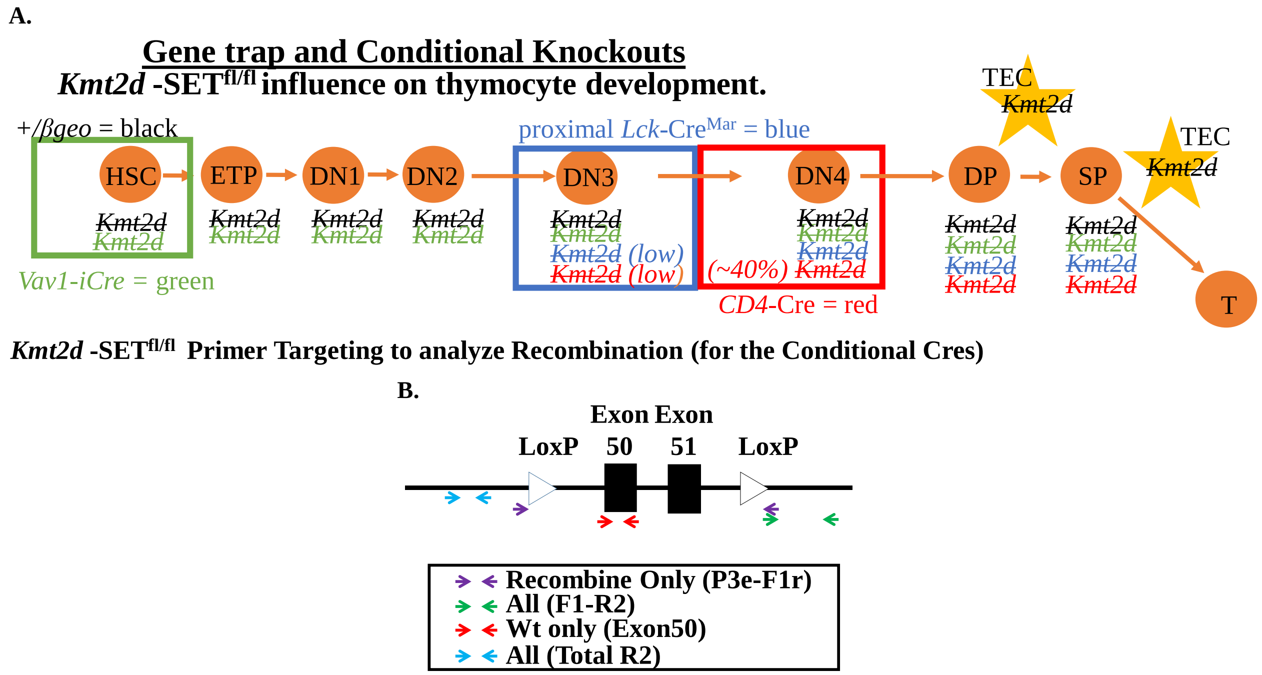


**Supplementary figure 2. Mouse models to analyze the role of Kmt2d during thymocyte/ T cell development through the use of differentially timed constitutional and conditional targeted *Kmt2d* loss.** (**a**) Conditional Cre system, when specific Cre recombinases turn on according to literature during thymocyte development depicted through loss of *Kmt2d*. Thymocytes in orange, supporting cells in yellow. Cres resulting in knockout *Kmt2d*: *Vav1*-iCre (green), *Lck*-Cre$^{Mar}$ (blue), *CD4*-Cre (red). Additionally, constitutive *Kmt2d*$^{+/\beta geo}$ mouse line shown in black (in both thymocytes and supporting cells). (**b**) Conditional mice harbor *LoxP* sites around *Kmt2d* exons 50 and 51 and disrupt the SET domain. Analysis of recombination qPCR primers targeted *Kmt2d* SET domain (Determination of recombination: purple, recombine only / green, all or the presence of Exon 50: red, wild-type only / blue all). See **Supplementary table 2** for primer sequence information and predicted primer outcomes.

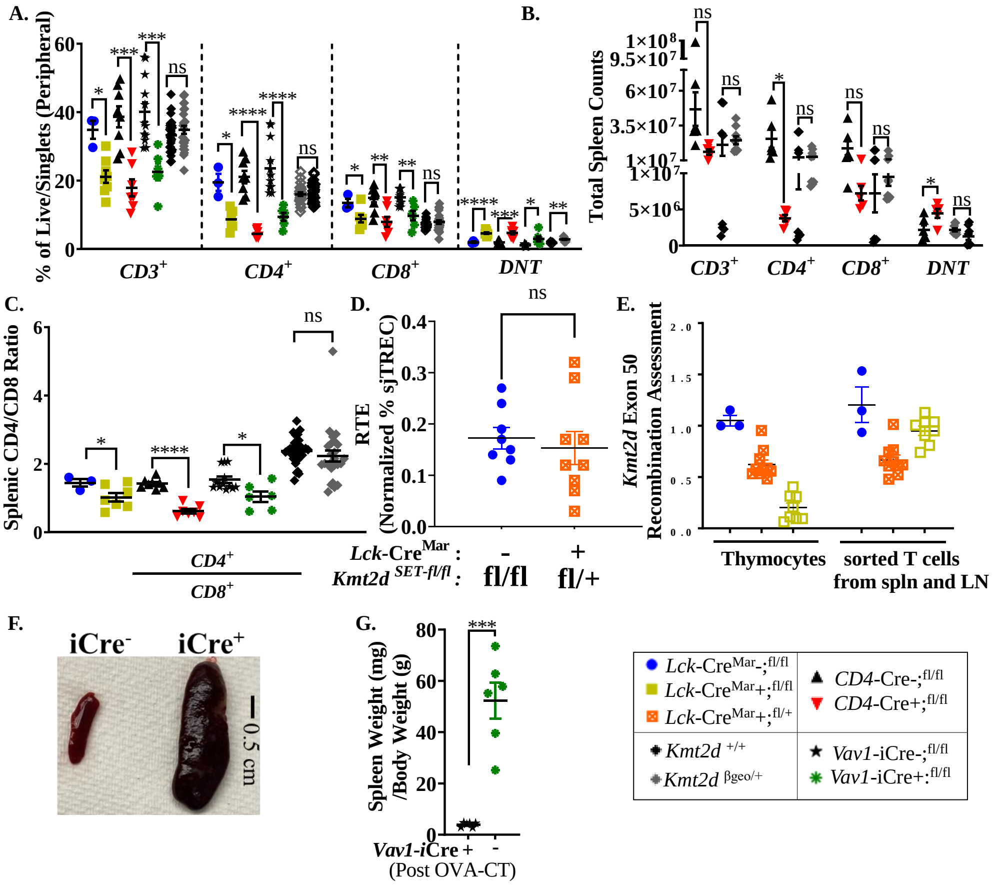


**Supplementary figure 3. *Kmt2d*-deficient peripheral T cells demonstrate decreased total T cells in live/single cells of spleen and altered CD4$^+$/CD8$^+$ ratios, as well as, analysis of heterozygous mice in RTE and recombination assays.** (**a**) Assessment of splenic T cell subpopulations as a percent of total live/singlets cells. (**b**) Total splenic cell counts of T cell populations and subpopulations. (**c**) Ratio of splenic CD4$^+$/CD8$^+$ in all mouse strains. (**d**) RTE in *Lck*-Cre$^{Mar}$ heterozygous-deficient mice (orange squares) compared to *Kmt2d*-sufficient littermates (blue circles) (sjTREC normalized to CD3$^+$). (**e**) Genomic DNA recombination assessment of exon 50 (SET domain), which is cleaved during Cre-recombination at *Loxp3* sites, compared to overall *Kmt2d* in unsorted thymocytes and bulk CD3$^+$ magnetic bead-sorted peripheral splenic/lymph node T cells of *Lck*-Cre$^{Mar}$ *Kmt2d*-homozygous knockout (yellow squares), *Kmt2d*-heterozygous deficient (orange squares), and control *Kmt2d*-sufficient littermates (blue circles). See schematic in **Supplementary figure 2**. (**f-g**) Example (**f**) of splenomegaly in sensitized *Vav1*-iCre-driven *Kmt2d* knockout and littermate controls and the quantified spleen weight normalized to total body weights (grams).

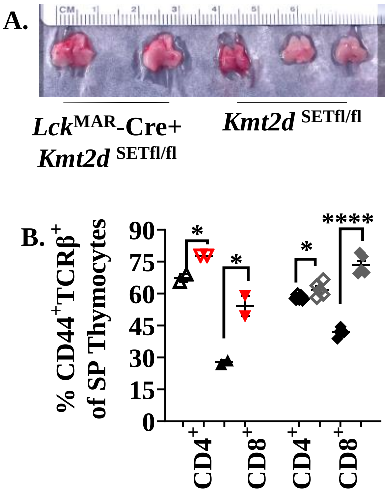


**Supplementary figure 4.** (**a**) Image of dissected *Lck*-Cre$^{Mar}$ *Kmt2d*-knockout thymus compared to littermate controls. (**b**) Cleavage of CD44$^+$ is required for SP to egress. Accumulation of CD44$^+$TCRβ$^+$ SP population (as a percent of CD4$^+$SP [open] or CD8$^+$SP [closed]) in *Kmt2d*-knockout (*CD4*-Cre; *Kmt2d*$^{\text{-SET-fl/fl}}$ [Cre$^-$ black/Cre$^+$ red triangles]) and *Kmt2d*-haploinsuffienct (*Kmt2d*$^{+/\beta geo}$ [$^{+/+}$ black/$^{+/\beta geo}$ grey diamonds]) cells. Significance labeled on graphs were determined using a parametric, unpaired, Welch's corrected *t-test* with *P*-values noted by stars based on *P*-value > 0.05 (ns), *P*-value < 0.05 (*) and *P*-value < 0.0001 (****).

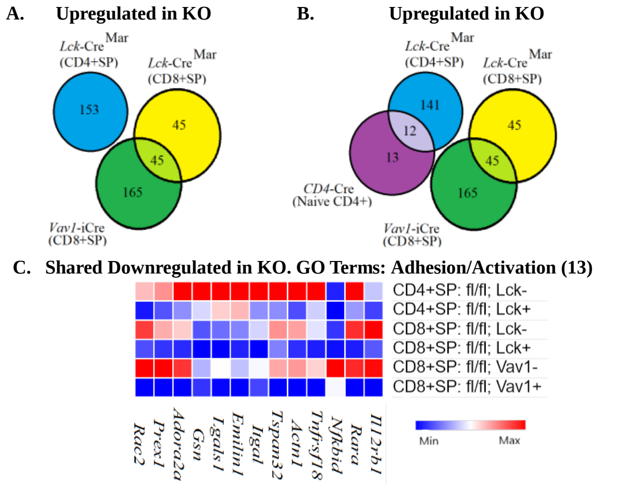


D. Gene location of Integrin Family

| Gene | Murine Chromosome # | Mouse Gene start/end | Human Gene |
|---|---|---|---|
| ***Itga4*** | chr2 | 79255425-79333114 | 2q31.3 |
| ***Itga6*** | chr2 | 71786938-71858427 | 2q31.1 |
| ***Itgae*** | chr11 | 73090582-73147449 | 17p13 |
| ***Itgal*** | chr7 | 127296259-127335137 | 16p11.2 |
| ***Itgb2*** | chr10 | 77530369-77565708 | 21q22.3 |
| ***Itgb7**** | chr15 | 102215994-102231935 | 12q13.13 |

*Note: Located Next to *Rarg*

E. Human integrin α / β Phylogenetic Trees

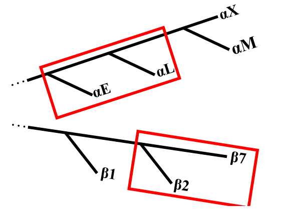


**Supplementary figure 5. Up-regulated expression analysis in *Kmt2d*-knockout mice.** (**a-b**) Potential *Kmt2d*- target genes estimated by selection of down-regulated genes in the *Kmt2d*-knockout compared to controls as *Kmt2d* normally regulates gene expression; therefore, these data reflect either indirect/downstream (negatively regulated) and/or bystander genes. Venn diagram demonstrates overlapping *Kmt2d*-knockout up-regulated filtered gene lists (Log2 fold change ≤ -0.59; adjusted *P*-value ≤ 0.1, RPKM ≥ 5 in at least one condition) between *Kmt2d* CD8$^{+}$SP (**a**) and CD8$^{+}$SP/CD4$^{+}$SP (**b**) as indicated from RNA-seq DESeq2 from *Vav1*-iCre [green] and *Lck*-Cre$^{Mar}$ [yellow] knockout CD8$^{+}$SP cells, CD4$^{+}$SP *Lck*-Cre$^{Mar}$ knockouts [blue], and peripheral naïve CD4$^{+}$ T cells *CD4*-Cre knockouts, GEO accession: GSE69162 (11) [purple]. Middle green oval/triangle are overlapping 45 (**a**,**b**) genes reflect shared up-regulated genes in CD8$^{+}$SP cells. Interestingly, neither the CD4$^{+}$SP, nor the peripheral CD4$^{+}$ cells overlap in up-regulated genes. (**c**) Morpheus heat map displays the average RPKM of 13 genes found within the shared GO: adhesion/activation molecules categories (top hits) from down-regulated 99 genes. Gradient color expression display, where blue (lower expression) titrates to red (higher expression): mouse strain by rows, genes are represented via columns. (**d**) Down-regulated integrins and their murine and human chromosomal locations. No observed clustered co-localization of down-regulated leukocyte-specific receptor integrin genes on both mouse and human chromosomes, but similar location of *Itga4* and *Itga6*. (**e**) Red box highlighting the proximity of the leukocyte-specific receptor integrins on their localized regions of their respective separate phylogenetic tree regions.

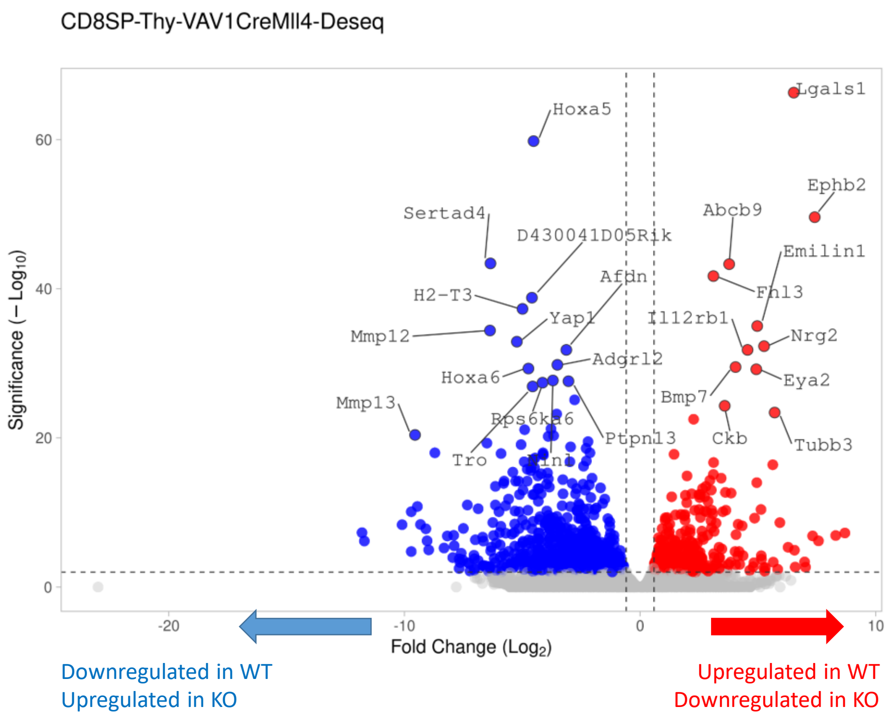


**Supplementary figure 6. Differential gene expression between CD8$^{+}$SP cells from conditional *Vav1*-iCre-driven *Kmt2d*-knockout and control littermates.** DESeq2 analyzed CD8$^{+}$SP RNA-seq from *Vav1*-iCre-driven *Kmt2d*-knockout and control littermates. Filtering Log2 fold change ≥ 0.59 (up-regulated in controls) or Log2 fold change ≤ $^{-}$0.59 (down-regulated in controls) and adjusted *P*-value ≤ 0.1 are depicted via dotted lines and filtered populations are displayed as down-regulation in the controls (as compared to the knockout [blue]) and up-regulation in controls (as compared to knockout [red]). Top 25 genes labeled with gene abbreviation.

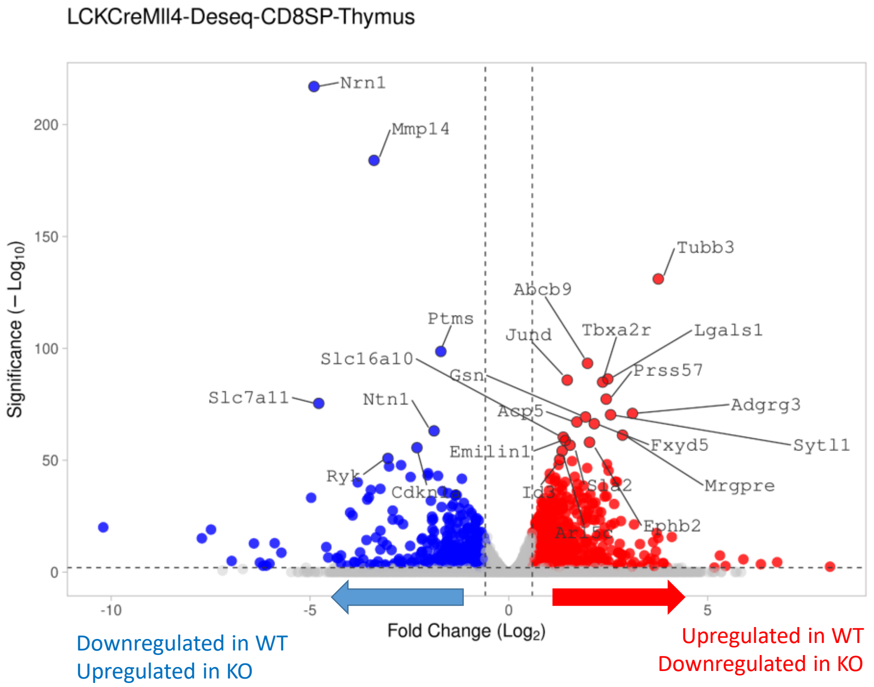


**Supplementary figure 7. Differential gene expression between $CD8^+$SP cells from conditional *Lck*-Cre$^{Mar}$-driven *Kmt2d*-knockout and control littermates.** DESeq2 analyzed $CD8^+$SP RNA-seq from *Lck*-Cre$^{Mar}$ -driven *Kmt2d*-knockout and control littermates. Filtering Log2 fold change ≥ 0.59 (up-regulated in controls) or Log2 fold change ≤ $^-$0.59 (down-regulated in controls) and adjusted *P*-value ≤ 0.1 are depicted via dotted lines and filtered populations are displayed as down-regulation in the controls (as compared to the knockout [blue]) and up-regulation in controls (as compared to knockout [red]). Top 25 genes labeled with gene abbreviation.

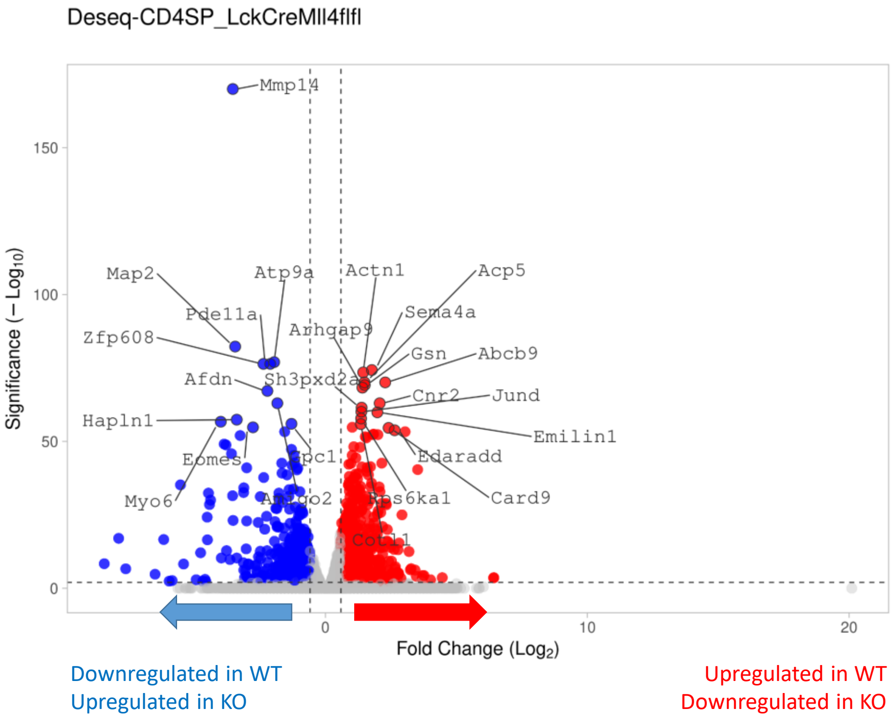


**Supplementary figure 8. Differential gene expression between $CD4^+$SP cells from conditional *Lck*-Cre$^{Mar}$-driven *Kmt2d*-knockout and control littermates.** DESeq2 analyzed $CD4^+$SP RNA-seq from *Lck*-Cre$^{Mar}$ -driven *Kmt2d*-knockout and control littermates. Filtering Log2 fold change ≥ 0.59 (up-regulated in controls) or Log2 fold change ≤ $^-$0.59 (down-regulated in controls) and adjusted *P*-value ≤ 0.1 are depicted via dotted lines and filtered populations are displayed as down-regulation in the controls (as compared to the knockout [blue]) and up-regulation in controls (as compared to knockout [red]). Top 25 genes labeled with gene abbreviation.

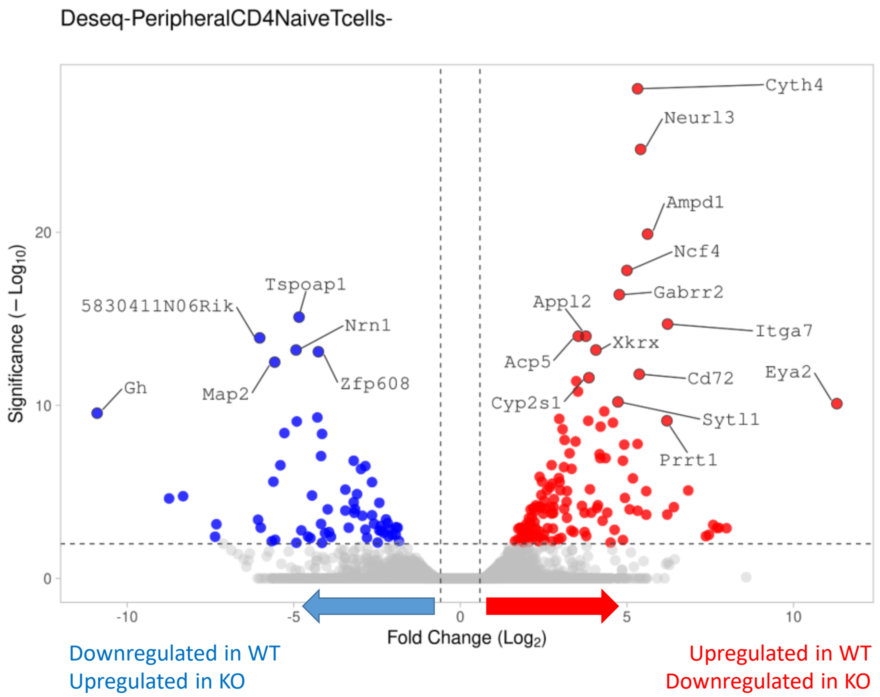


**Supplementary figure 9. Differential gene expression between peripheral naïve CD4$^{+}$ cells from conditional *CD4*-Cre-driven *Kmt2d*-knockout and control littermates.** DESeq2 analyzed peripheral naïve CD4$^{+}$ T cells RNA-seq from GEO accession: GSE69162 (11) *Kmt2d*-knockout and control littermates. Filtering Log2 fold change ≥ 0.59 (up-regulated in controls) or Log2 fold change ≤ $^{-}$0.59 (down-regulated in controls) and adjusted *P*-value ≤ 0.1 are depicted via dotted lines and filtered populations are displayed as down-regulation in the controls (as compared to the knockout [blue]) and up-regulation in controls (as compared to knockout [red]). Top 25 genes labeled with gene abbreviation.

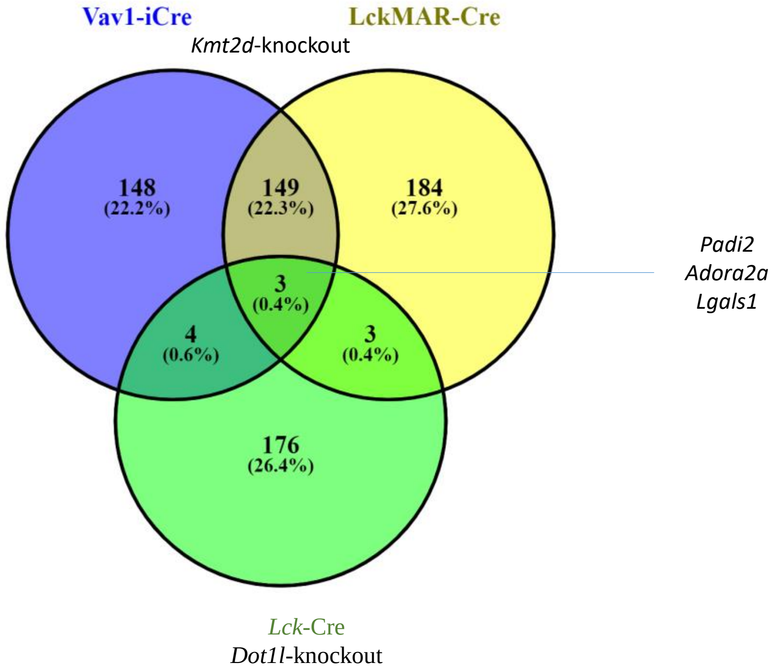


**Supplementary figure 10. Overlapping differential gene expression between CD8$^+$SP cells from conditional *Vav1*-iCre- and *Lck*-Cre$^{Mar}$-driven *Kmt2d*-knockout and *Dot1l*-knockout mice.** Venn diagram demonstrates overlapping *Kmt2d*-knockout down-regulated filtered gene lists (Log2 fold change ≥ 0.59; adjusted *P*-value ≤ 0.1; RPKM > 5 in at least one condition) between CD8$^+$SP *Kmt2d*–knockout (*Vav1*-iCre [green] and *Lck*-Cre$^{Mar}$ [yellow]) and *Dot1l*-knockout mice (*Lck*-Cre$^{Mar}$ [blue] from GEO accession: GSE138910 (60)) from RNA-seq generated DESeq2. Middle green oval/triangle are overlapping three genes reflect shared down-regulated genes in all CD8$^+$SP cells.

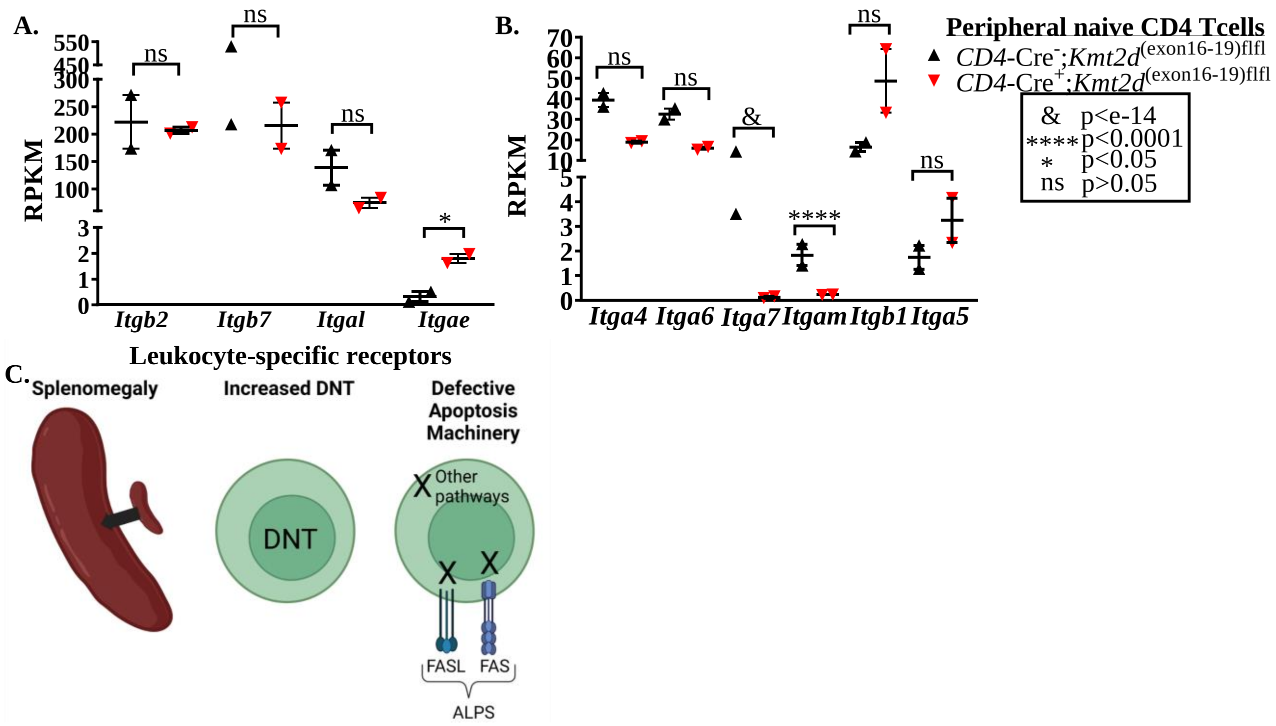


**Supplementary figure 11. Analysis of integrins in peripheral naïve $CD4^+$ T cells.** (**a-b**) RPKM of leukocyte-specific receptor (**a**) integrin and non-leukocyte specific (**b**) integrins that are down-regulated in *Kmt2d* knockouts and up-regulated non-leukocyte specific integrins from peripheral naïve $CD4^+$ T cells from *CD4*-Cre; *Kmt2d* $^{exon\ 16\text{-}19\text{-}fl/fl}$ ($Cre^-$ black up-right triangles/$Cre^+$ red inverted triangles). Each dataset displays black lines, which represent mean ± SEM. (**c**) The ALPS defining characteristics (splenomegaly, increased percentage of double-negative T cells, and defective apoptotic machinery). Created with BioRender.com. Significance (adjusted *P*-values are generated from DESeq2 analysis. *P*-value > 0.05 (ns), *P*-value < 0.0001 (****), and *P*-value < $1e^{-14}$ (&).

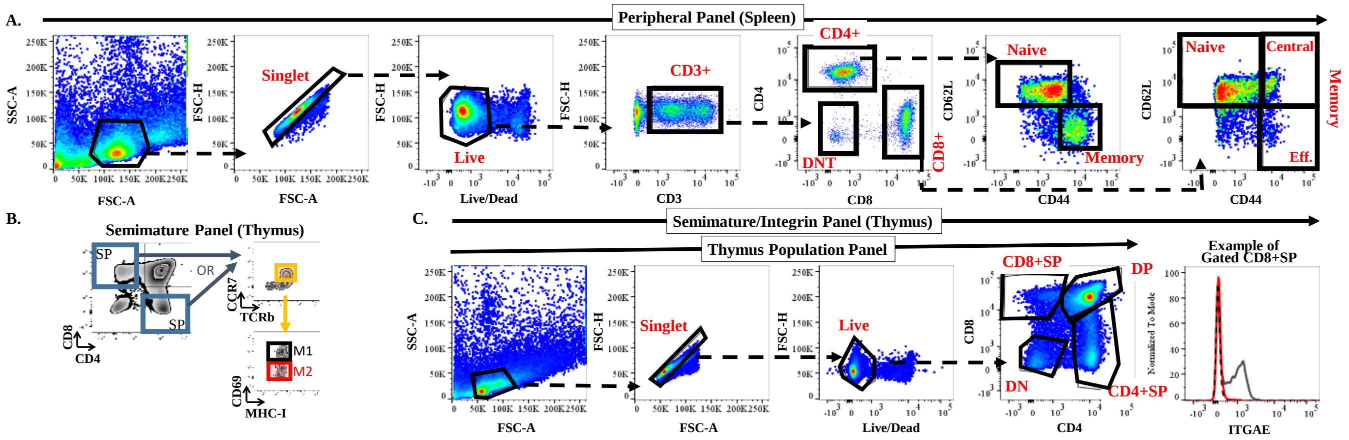


**Supplementary figure 12. Gating Strategy of thymocyte populations.** (**a**) Peripheral Panel (of splenocytes): First, gate on size/granularity (FSC-A vs. SSC-A); on singlets (FSC-A vs. FSC-H); on live (Live/Dead stain negative vs. FSC-H), on CD3 (CD3 positive vs. FSC-H). Next, gate on CD4 by CD8 to separate out the CD4+, CD8+ and the CD4-CD8- (double-negative, DNT) populations. Finally, gate on CD44 by CD62L to separate naïve (CD62L$^{hi}$) versus memory (CD44$^{hi}$) for CD4+. For CD8+, additionally determination of effector memory (CD62Llo) compared to central memory (CD62L$^{hi}$) was defined by expression of CD62L levels on CD44$^{hi}$ cells. (**b**) Semimature Panel (for the Thymus). Similarly to (**a**), gate on CD4 by CD8 to separate out the CD4+single-positive (SP) and CD8+SP populations. Each SP population gating of TCRβ by CCR7 was used to gate on CCR7$^{hi}$ cells. Lastly, M1 compared to M2 was determined through CD69 expression (as co-gated on MHCI). (**c**) Semimature/Integrin Panel (Thymus). Similar gating strategy of size/granularity, singlet, and live cells as depicted in the peripheral gating (**a**), with subsequent gating on CD4 and CD8 to determine double-negative (DN), double-positive (DP) and single positive (CD4$^{+}$SP, CD8$^{+}$SP) thymocytes. Each population was assessed for integrin expression and intensity.
